# Quantifying eco-evolutionary contributions to trait divergence in spatially structured systems

**DOI:** 10.1101/677526

**Authors:** Lynn Govaert, Jelena H. Pantel, Luc De Meester

## Abstract

Ecological and evolutionary processes can occur at similar time scales, and hence influence one another. There has been much progress in the development of metrics that quantify contributions of ecological and evolutionary components to trait change over time. However, many empirical evolutionary ecology studies document genetic differentiation among populations structured in space. In both time and space, the observed differentiation in trait values among populations and communities can be the result of interactions between non-evolutionary (phenotypic plasticity, changes in the relative abundance of species) and evolutionary (genetic differentiation among populations) processes. However, the tools developed so far to quantify ecological and evolutionary contributions to trait change are implicitly addressing temporal dynamics because they require directionality of change from an ancestral to a derived state. Identifying directionality from one site to another in spatial studies of eco-evolutionary dynamics is not always possible and often not desired. We here suggest three modifications to existing metrics so they allow the partitioning of ecological and evolutionary contributions to changes in population and community trait values across landscapes. Applying these spatially modified metrics to published empirical examples shows how these metrics can be used to generate new empirical insights and to facilitate future comparative analyses. The possibility to apply eco-evolutionary partitioning metrics to populations and communities in real landscapes is critical as it will broaden our capacity to quantify eco-evolutionary interactions as they occur in nature.

## Introduction

During the past decade ecologists and evolutionary biologists have become increasingly aware that ecological and evolutionary processes can combine to structure populations and communities (Hairston et al. 2005; Schoener 2011; Barraclough 2015; Hendry 2017). This has prompted development of a suite of metrics to describe and quantify eco-evolutionary contributions to numerous processes that were traditionally considered to result only from ecological dynamics (Hairston et al. 2005; Collins and Gardner 2009; Ellner et al. 2011; Govaert et al. 2016). These methods, however, have generally been developed for and applied to populations and communities separated in time. For example, a study by Becks et al. (2012) used experimental chemostats to show that over a time period of 90 days evolutionary responses in the defense traits of an algal prey were more important to rotifer population growth than changes in algal abundance. To come to these conclusions they used an eco-evolutionary partitioning metric developed by Ellner et al. (2011). In another study, the same metric was used to compare the relative impact of an environmental change (dry or wet soil) and of plant evolutionary history (adaptation during 16 months to a dry or wet environment) on soil microbial and fungal community diversity (terHorst et al. 2014). They found that rapid evolutionary responses of plant populations to drought are as important as the direct ecological effects of this stressor, indicating that ecological and evolutionary effects had similar magnitudes. Gómez et al. (2016) similarly found that pre-adaptation to elevated temperature for 48 days in *Pseudomonas fluorescens* contributed as much to change in taxon composition of a compost bacterial community as the presence of the species *P. fluorescens* itself.

While eco-evolutionary dynamic studies are mainly focused on reciprocal evolutionary and ecological changes over time, many evolutionary ecology studies of natural systems consider trait variation among geographically segregated patches and trait turnover along particular spatial gradients rather than over time. Spatial landscape heterogeneity plays a role in shaping genetic structure in natural populations (Ackerman et al. 2013), and the distribution of phenotypes of local populations in space exposed to different environments may diverge through adaptive plasticity and local adaptation (Via and Lande 1985; Kawecki and Ebert 2004; Logan et al. 2016). Similarly, species sorting across the landscape might also occur when different environments are dominated by different competing species (Fox and Harder 2015). Therefore, spatially separated populations or communities may be structured by non-evolutionary (i.e. phenotypic plasticity and species sorting) and evolutionary processes. Quantifying these processes in spatial study systems may improve our understanding on how geographical, environmental and community features structure evolutionary and non-evolutionary contributions to trait variation in populations and communities inhabiting natural landscapes.

As evolutionary trait change occurs over time, studies quantifying trait evolution often have a clear direction in the observed trait change from past to present. However, studies measuring trait divergence among spatially separated populations or communities capture the trait values of the species at a defined moment in time. While the observed trait divergence in these studies does reflect past evolutionary changes, information about past states is often not available. Whether all populations diverged from a common ancestor simultaneously or fragmented at various points throughout the past is unknown. In such instances, the goal may be to quantify the amount of trait divergence that can be attributed to evolutionary and non-evolutionary contributions, rather than to determine such contributions to trait change assuming a certain ancestry. However, some spatial study systems do imply a directed trait change among spatially separated populations or communities. For example, the direction of ancestry is known in studies that investigate invasion history (e.g. stickleback colonisation of freshwater lakes; Bell et al. 2004; Le Rouzic et al. 2011) or that look at range expansions where the peripheral population can be traced back to more central populations (e.g. Safriel et al. 1994; Volis et al. 2001; Swaegers et al. 2014). In some studies, ancestor-descendant relationships are assumed (e.g. Etterson and Shaw 2001). In these studies, eco-evolutionary partitioning metrics that quantify non-evolutionary and evolutionary components of temporal trait change can be used without modification.

To our knowledge, no study to date has partitioned population or community structure in a spatially explicit scenario, even though spatial variation in traits, genes, and species composition across landscapes are central to studies that focus on ecological or evolutionary processes in isolation. The study of Norberg et al. (2012) considers spatially structured communities in a spatially explicit model, but still quantifies ecological and evolutionary contributions to temporal community trait change as opposed to spatial community trait divergence. The study of Pantel et al. (2015) modified the metric proposed by Hairston et al. (2005) and Ellner et al. (2011) to assess genetic and environmental contributions to measures of community composition (e.g. species richness, change in Simpson’s diversity), and their experimental question and methods did not utilize a direction of change between communities. However, they did not explicitly quantify among-community trait variation in the contribution of evolutionary and ecological components. Questions such as whether the magnitude or relative importance of non-evolutionary and evolutionary effects is related to features of the organisms (e.g. generation time) or the landscape (e.g. degree of isolation or degree of habitat heterogeneity), or when populations and communities are more likely to respond to environmental change via shifts in the relative abundances of species, phenotypic plasticity, evolutionary change, or some combination of these is an important next step for studies of eco-evolutionary dynamics. However, in order to answer these questions, we need appropriate tools to quantify non-evolutionary and evolutionary contributions to population and community trait divergence among landscapes or experimental units.

Today, a handful of metrics is available to calculate non-evolutionary and evolutionary contributions to trait change such as the Price equation (Price 1970; Price 1972), metrics based on reaction norms (Hairston et al. 2005; Ellner et al. 2011; Govaert et al. 2016) and the recently developed Price-Reaction-Norm equation (Govaert et al. 2016). These metrics have previously been compared to one another, with differences in their set of assumptions highlighted (Govaert et al. 2016; Govaert 2018) and thus can provide a first step to explore modifications to spatial study systems. The goal of this study is to extend these eco-evolutionary partitioning metrics to assess how non-evolutionary and evolutionary processes, and their interactions, contribute to observed trait differentiation between spatially structured populations and communities. Specifically, our study will address the large number of studies that do not seek to understand a change across a direction, but instead compare differentiation among groups. We develop modifications to existing partitioning metric for temporal data and show that depending on these modifications, the assessment of the importance or the interpretation of the contributing ecological and evolutionary processes may change. We then apply the modified metrics to selected empirical datasets to illustrate the diversity of research questions that can be addressed and how the here developed metrics can be used to facilitate comparative analyses and generate new empirical insights.

### Applying current metrics to spatially structured systems

Spatially separated populations and communities are structured by ecological and evolutionary processes that may result in trait differentiation among these populations and communities (Via and Lande 1985; Kawecki and Ebert 2004; Fox and Harder 2015; Logan et al. 2016). In order to quantify the relative importance of evolution as opposed to ecology in structuring spatial populations and communities, it is important to separate evolutionary change (at the species level) from phenotypic plasticity (at the individual level), and from ecological change (i.e. species sorting; at the community level). Each of these components contributes to phenotypic community trait change, and their effects are likely not independent from one another, implying that their interactions (e.g. evolution of plasticity, species sorting × evolution) should be taken into account as well. This general separation of components at different organisational levels is referred to as the ‘eco-evo sandwich’, where evolutionary change at the intermediate (species) organizational level can interact with ecological processes at lower (individual) and higher (community) organizational levels (Fig. 1; modified from Govaert et al. 2016).

**Figure 1:**
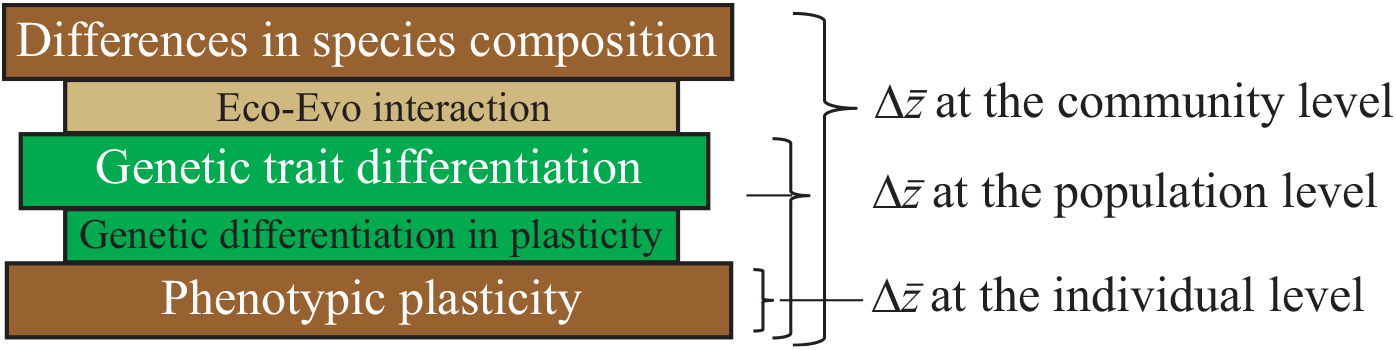
Visualisation of the ‘eco-evo sandwich’ modified from Govaert et al. (2016) to a spatial study. Community trait divergence can be separated in plasticity effects at the individual level (brown bar), genetic trait differentiation at the population level (green bar), differences in species composition at the community level (i.e. species sorting; brown bar) and interactions between these non-evolutionary and evolutionary processes. These interactions may involve genetic differentiation in plasticity (smaller green bar) which is an evolutionary process, and a species sorting × evolution interaction (smaller light brown bar).

Most currently used eco-evolutionary partitioning metrics depend on the choice of the reference point (Fig. 2A). Hence, the direction of change might alter the result of existing eco-evolutionary partitioning metrics that depend on direction. For example, applying current metrics such as the Price equation or reaction norm approach to partition a trait shift between two hypothetical populations of a single species shows that depending on the reference population chosen (i.e. partitioning trait shift from Population 1 to 2 or from Population 2 to 1) a different contribution of evolution is found (Fig. 3). This difference is an undesirable consequence of applying eco-evolutionary partitioning metrics that are designed to separate directional trait change to populations or communities for which there is no information on their ancestry. We therefore suggest three approaches, illustrated in Figure 2B, to modify current partitioning metrics to quantify ecological and evolutionary processes to undirected trait shifts, typically assessed in spatial study systems. In subsequent sections, we provide formulae as well as demonstrative calculations for these modifications. We then apply these approaches to three empirical studies.

**Figure 2:**
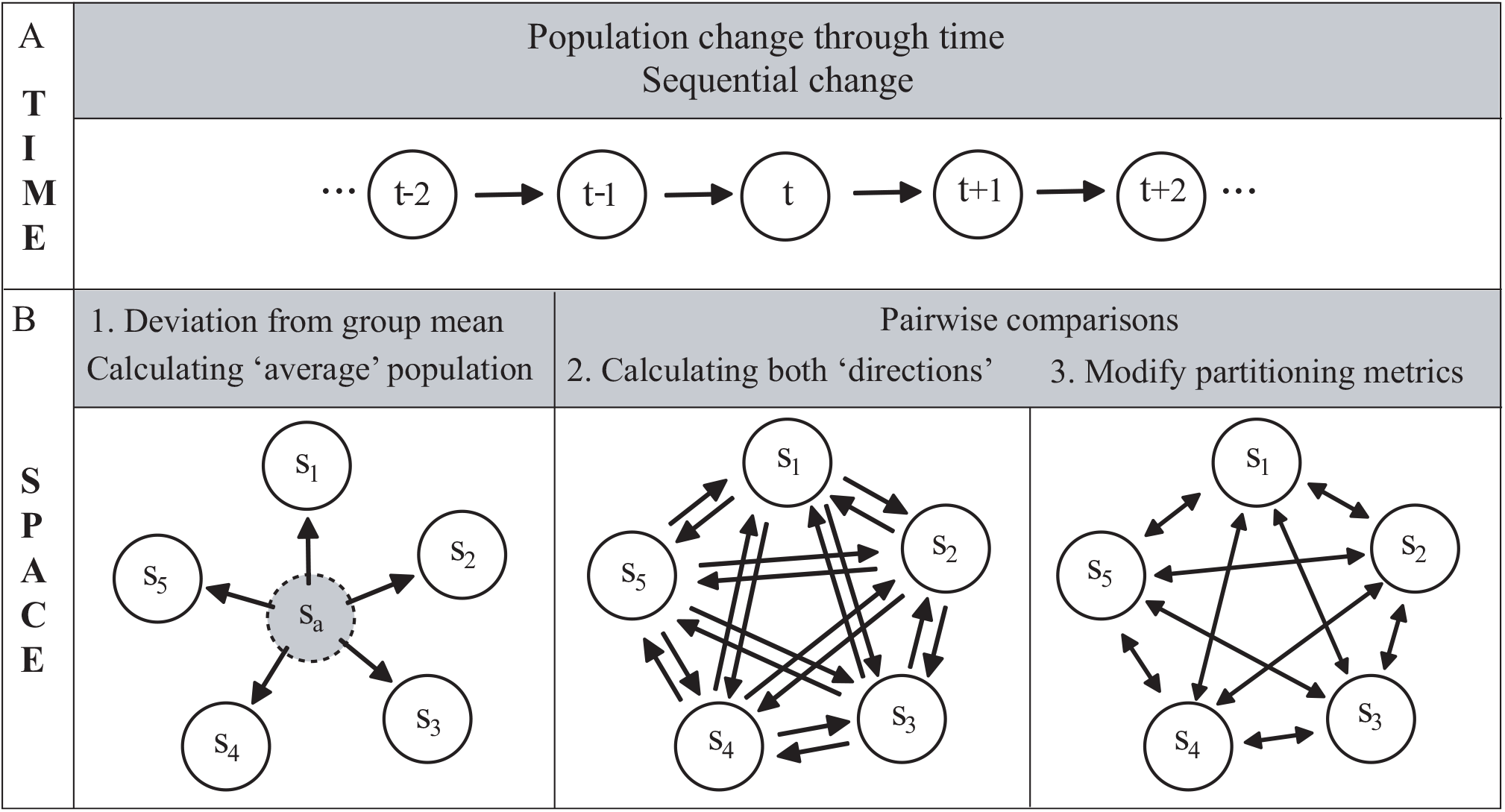
Visual representation of (A) directed trait change in a temporal study system where trait change goes from time *t* − 1 to *t* to *t* + 1 etc. and (B) trait divergence in spatial study systems here consisting of 5 sites, where non-evolutionary and evolutionary contributions to trait divergence can be estimated either as deviations from a common group mean or known ancestor (Approach 1) or as an averaged change between pairs of sites by calculating the non-evolutionary and evolutionary contributions from e.g. site *s*_1_ to site *s*_2_ and vice versa followed by averaging the absolute values of those quantities and this for all pairwise combinations (Approach 2) or by using partitioning metrics modified to undirected trait change (Approach 3). Arrows do not reflect spatial distances, but directions of trait change to which non-evolutionary and evolutionary contributions are calculated for.

**Figure 3:**
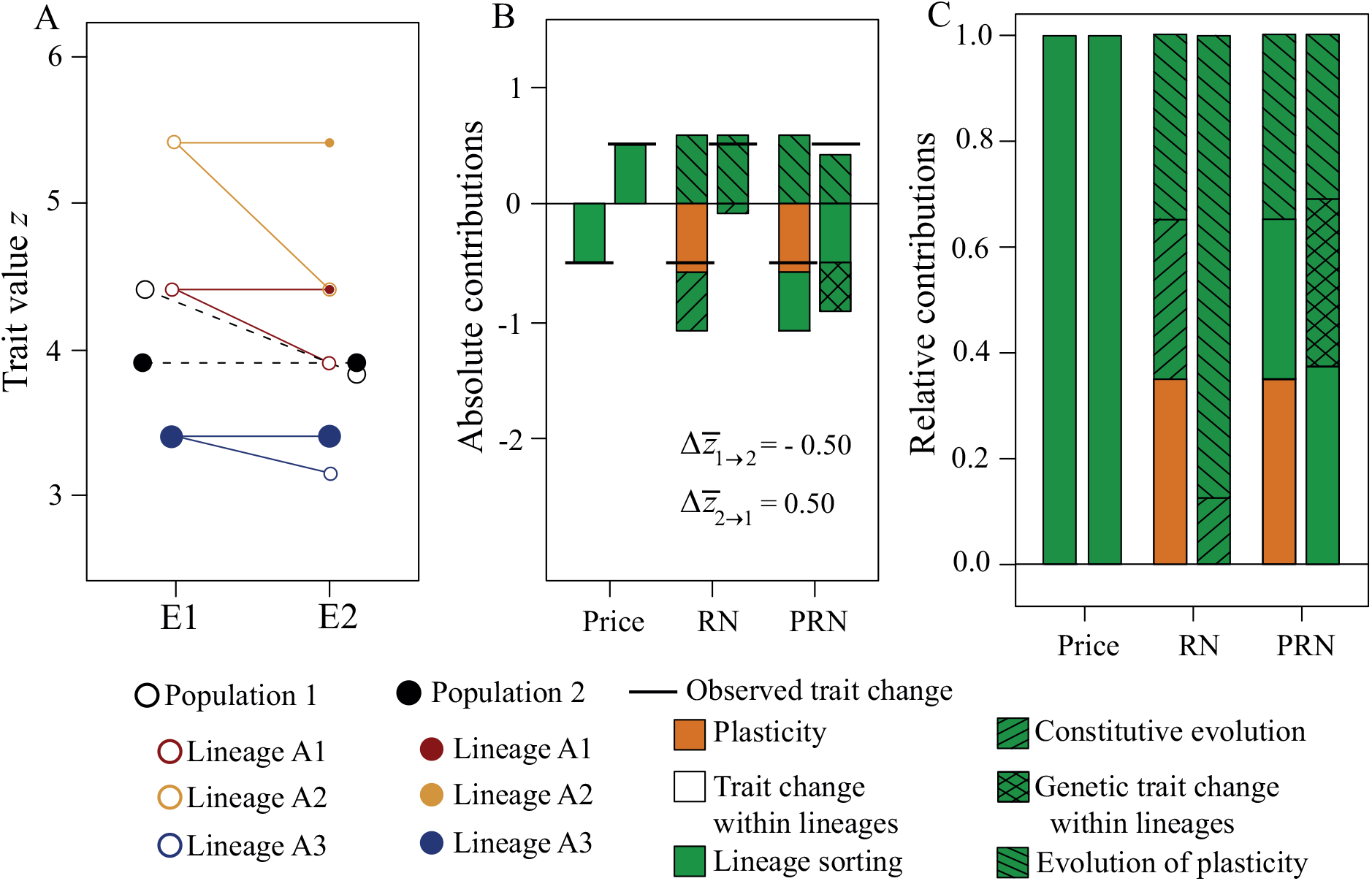
Numerical example displaying shift in a trait *z* between Population 1 at site 1 (unfilled black circles) and Population 2 at site 2 (filled black circles) in which the populations consist of the same three genetic lineages (indicated by blue, red, yellow color). The populations differ in environmental condition E, i.e. Population 1 (resp. 2) originated from an environmental condition E1 (resp. E2). Individuals of the three lineages were measured for trait *z* in both environmental conditions in order to construct reaction norms. (A) Visualisation of the reaction norms of the three lineages, where the unfilled (resp. filled) circles represent the average trait value *z* of the genetic lineage at site 1 (resp. 2) at environmental condition E1 and E2. The black unfilled (resp. filled) circles represent the average trait value of the total population calculated as a lineage abundance-weighted mean using the lineage trait values and lineage relative abundances (given by the size of the symbols). (B) Absolute and (C) relative contributions of the evolutionary and non-evolutionary components to the average trait shift from Population 1 to Population 2 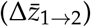 and vice versa 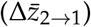 using the Price equation (Price), the reaction norm approach (RN) and the Price-Reaction-Norm equation (PRN). In (B) the vertical bar represents the observed trait difference between Population1 at E1 and Population 2 at E2, either calculated as 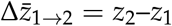 or 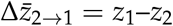.

A first modification involves setting a common reference by constructing a midpoint such as the group mean. This is conceptually similar to using Helmert contrasts in regression analysis. This way we partition trait deviation for each site from the group mean into non-evolutionary and evolutionary processes (Fig. 2B). In the second modification, we quantify non-evolutionary and evolutionary contributions to directed trait change of both directions (treating each of the two sites in each pair as a reference), and then average the resulting fractions (Fig. 2B). In a third modification, we explore ways to mathematically adapt current eco-evolutionary partitioning metrics to become independent of the reference chosen. The metrics we propose can be applied to both population as well as community trait data, but we focus on the metrics applied to populations in the main text and provide their extension to community data in Appendix A.

### Price, reaction norm and Price-Reaction-Norm as applied to populations

Because our intention is to modify the metrics for non-directional comparisons, we first present the basic equations for the Price equation and the Reaction Norm approach for directional comparisons (usually for two points in time). We exclude the Price-Reaction Norm approach from the main text as this is a combination of both the Price and Reaction Norm equation, but provide the details for that equation in Appendix B.

The Price equation, introduced by G. R. Price (Price 1970; Price 1972), has been used to describe trait change in a biological population from one generation to the next. The Price equation is very versatile and has proven its usefulness in evolutionary biology (detailed in Queller 2017), ecology (Fox 2006; Fox and Kerr 2012), epidemiology (Day and Gandon 2006), and evolutionary ecology (Collins and Gardner 2009; Ellner et al. 2011; Govaert et al. 2016). We here use the version of the Price equation that partitions trait change between two time points in an asexually reproducing population consisting of *N* genetic lineages, uniquely indexed by *j* ∈ {1, … , *N*} (Govaert et al. 2016). For this metric to apply, we require information on the relative abundance and average trait value for each lineage in each population at both time points. This metric then divides trait change between two time points into a component that gives the changes in the relative abundances of the lineages (i.e. lineage sorting) and a component that gives the trait change within lineages:

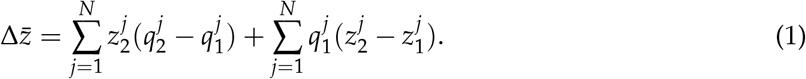

In this equation, 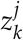 (resp. 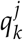) represents the trait value (resp. relative abundance) of lineage *j* at time *t*_*k*_. There exist many organisms that reproduce asexually and for which the Price equation might be suitable. For example, clonal plants (e.g. *Arabidopsis thaliana*, *Solidago altissima*, North American *Taraxacum officinale*) and seagrasses (e.g. *Zostera marina*), or crustaceans that undergo obligate parthenogenesis (e.g. some morphs of *Daphnia pulex*) are some of the many species for which distinct genetic lineages might differentially assemble into populations and contribute to trait shifts.

The reaction norm approach uses the concept of reaction norms originally introduced by R. Woltereck (‘*Reaktionsnorm*’; Woltereck 1909). Reaction norms have been widely used in quantitative genetics to determine genotype-by-environment interactions. A reaction norm gives a formal association between a phenotype, its genotype and the environment, by mapping each genotype onto its phenotype as a function of the environment (Stearns 1989). Stoks et al. (2016) and Govaert et al. (2016) used mean reaction norms of a population (i.e. average reaction norm across all individuals or genotypes of that population) to assess the contributions of ancestral plasticity, constitutive evolution and evolution of plasticity to population trait change between two time points. This metric uses population means, and partitions the observed trait change between two time points as follows:

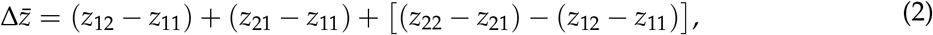

where *z*_*kl*_ is the average trait value of the population at genetic state *k* (i.e. sampled at time point *t*_*k*_) in environmental condition *l*. This metric thus requires information on the trait values of representative sets of individuals of both populations in the two environmental conditions that correspond to the habitat each population is found in. The first term on the right hand side of eqn (2) reflects phenotypic plasticity, the second term reflects constitutive evolution, and the last term reflects evolution of plasticity.

### Spatial modification of the Price and reaction norm equation

#### Approach 1: Deviation from group mean

Contrasts in regression models determine how the model coefficients of categorical variables are interpreted. Treatment contrasts, for example, sets a baseline (reference) and compares all subsequent levels to this baseline. Depending on the baseline, interpretation of the model coefficients can change. Helmert contrasts compare each level against a mean of the preceding levels, and scaling the contrasts can allow comparing two binary treatment levels to their midpoint (average). Eco-evolutionary partitioning among spatially separated populations can in a similar way be performed by using the same reference for comparisons with all observed populations by constructing a group mean that serves as the baseline. Current eco-evolutionary partitioning metrics can then still be used to calculate deviations from this midpoint, i.e. the group mean, for each observed population. Constructing the group mean can differ depending on the type of metric used, and we therefore show how this mean can be constructed when using data applicable for the Price equation or the reaction norm approach (Price-Reaction-Norm equation detailed in Appendix B).

##### The Price equation

We here assume that each population consists of the same *N* genetic lineages from which we have information on the relative abundances and trait values. While this assumption might be unlikely in natural landscapes consisting of spatially separated populations, it is used here to facilitate illustration of the procedure. A detailed explanation when this assumption is not met (i.e. when sites differ in the presence and absence of different lineages) is given in Appendix C. Constructing a group mean requires calculating average trait values 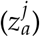 and average relative abundances 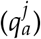 of the lineages across the populations, which are calculated as:

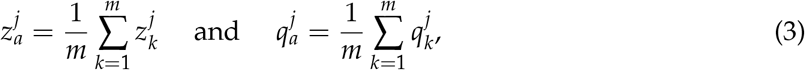

where 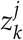 and 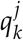 denotes the average trait value and relative abundance of lineage *j* of Population *k*. Using the Price equation, we can partition the observed population trait deviation of Population *k* from the group mean as follows:

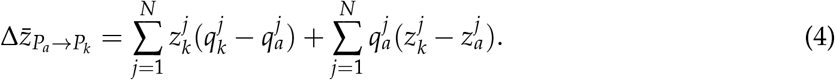

The first term in the right hand side of eqn (4) refers to the trait deviation due to lineage sorting and the second term refers to the trait deviation due to within-lineage trait deviation.

##### The reaction norm approach

The construction of the group mean for the reaction norm approach can be seen as an average population assumed to originate from an average environmental condition (see Appendix D for a graphical explanation of creating this group mean). In this case we assume that the set of *m* populations can be subdivided into two groups based on an environmental conditions (e.g. populations in which a specific predator is present or absent or populations experiencing high or low nutrients). In order to construct this group mean we need to calculate *z*_*al*_ (i.e. the average trait value of the average population in environmental condition *l* ∈ {1, 2}), *z*_*ka*_ (i.e. the average trait value of Population *k* in the average environmental condition *a*), and *z*_*aa*_ (i.e. the average trait value of the average population in the average environmental condition *a*) as follows:

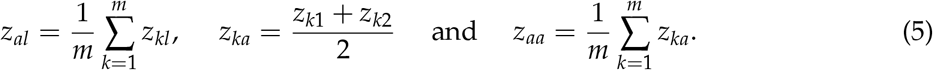

The reaction norm equation can then be used to partition the observed trait deviation of Population *k* from the average population as:

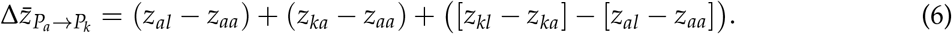

The first term on the right hand side of eqn (6) is the plasticity component, the second term is the trait deviation due to genetic trait differentiation, and the third term is the trait deviation due to a genetic differentiation in plasticity. It is important to keep in mind that the plasticity component does not reflect the absolute amount of plasticity but rather reflects the average phenotypic plasticity response. Note that as the last component on the right hand side of eqn (6) approaches zero for a certain Population *k*, the more similar degree of plasticity the population has with the group mean. The greater this value is, the more genetically differentiated the population is in its plasticity response compared to the group mean.

#### Approach 2: Average of components in both directions

Calculating eco-evolutionary deviations of each observed population from a group mean provides insight into the variation in non-evolutionary and evolutionary contributions among the set of populations. However, one may also compare pairs of populations and then summarise the observed non-evolutionary and evolutionary contributions among all pairs of sites to get an overall magnitude effect of evolution and ecology. We therefore propose a second modification to determine the relative contribution of different eco-evolutionary processes to shifts in traits among populations that treats each population as a reference. Consider for example two populations inhabiting spatially distinct sites. One could calculate non-evolutionary and evolutionary components to the trait divergence (i.e. 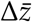) using first the population at site 1 as a reference, and subsequently using the population at site 2 as a reference. Absolute values of the non-evolutionary and evolutionary components obtained from both calculations are then averaged to quantify the overall relative importance of evolution and ecology. We here formulate how this approach translates to the two types of eco-evolutionary partitioning metrics between pairs of spatially separated populations (in what follows the populations used in the pairwise comparison are stated as Population 1 and Population 2).

##### The Price equation

The Price equation given by eqn (1), assumes Population 1 to be the reference, and partitions the observed trait shift from Population 1 to Population 2 for a trait *z* (i.e. 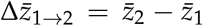) into evolutionary and non-evolutionary contributions. Using Population 2 as a reference and partitioning a trait shift from Population 2 to Population 1, using the Price equation, partitions an observed trait difference 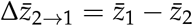 as follows:

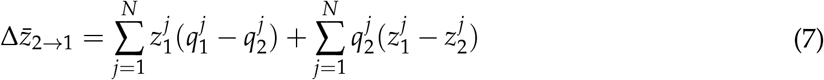

The first term on the right hand side of equation (1) and (7) gives the change in the relative abundances of the lineages, either 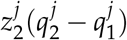 in eqn (1) or 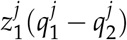 in eqn (7). These two terms only differ in the trait value that is multiplied with the change in relative abundances of the lineages, which is either 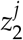 in eqn (1) or 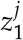 in eqn (7). Averaging the absolute values of these two terms gives the overall magnitude of lineage sorting to the trait divergence between Population 1 and 2, i.e.

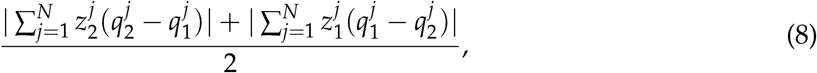

Similarly, one can calculate the overall magnitude of the trait difference within genetic lineages by averaging the absolute values of the two last terms in equation (1) and (7), i.e.

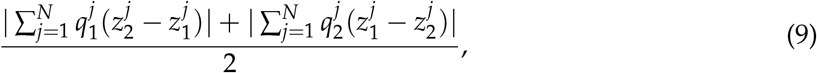

Dividing eqn (8) and (9) by their sum gives the overall relative importance of each process.

##### The reaction norm approach

The reaction norm approach given in eqn (2), partitions observed population trait change from one time point to the next into an ancestral plasticity, constitutive evolution and evolution of plasticity component. For two spatially separated populations, however, we can calculate the different components using either Population 1 or Population 2 as reference. Partitioning trait divergence from Population 1 to Population 2, hence using Population 1 as a reference, is given by eqn (2). When using Population 2 as a reference, the observed trait difference equals 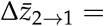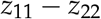 and can be partitioned into the following components:

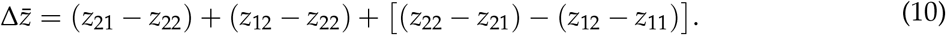

The first term on the right hand side of eqn (10) is the plasticity response of Population 2, the second term is the genetic trait differentiation from Population 2 to 1, and the last term gives the change in plasticity from Population 2 to 1. Averaging the absolute value of the plasticity components of eqn (2) and eqn (10) then gives the absolute magnitude of plasticity to the trait difference between Population 1 and 2:

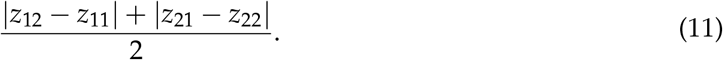

Similarly, averaging the genetic trait differentiation components and the genetic differentiation in plasticity components of eqn (2.2) and eqn (2.11) gives the absolute magnitude of genetic trait differentiation:

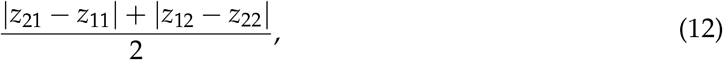

and of genetic differentiation in plasticity, respectively:

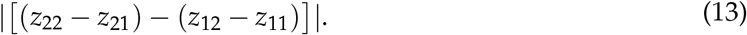

Note that the genetic differentiation in plasticity is the same value in both directions, so the average of the absolute values of genetic differentiation in plasticity in both directions equals the absolute value of one direction.

#### Approach 3: Partitioning metrics for undirected trait change

Approach 1 introduces a direction in the study system by constructing a group mean. This provides a common reference from which ecological and evolutionary contributions to trait deviations can be quantified. Approach 2 eliminates directionality by averaging the obtained contributions of ecology and evolution from the two possible directions of trait change between a pair of populations. Thus, both Approach 1 and 2 still use eco-evolutionary partitioning metrics for directed trait change. We here propose a third modification in which we explore ways to mathematically modify partitioning metrics to make them independent of a reference. The nature of this modification depends on the metric itself, and hence the interpretation of the resulting components also differs. The modification suggested here is used for pairwise comparison, and we denote the two populations used in the pairwise comparison as Population 1 and Population 2.

##### The Price equation

The Price equation can be used to separate trait change between populations into a lineage sorting and a within-lineage trait change component. Depending on whether we partition the trait change from Population 1 to Population 2 (as in eqn (1)) or vice versa (as in eqn (7)), the contribution of lineage sorting equals the difference in relative abundances of the lineages between both populations multiplied with the trait value of the lineages of either Population 1 (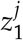) or Population 2 (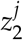). Multiplying this term instead with an average lineage trait value 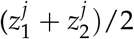 as opposed to the lineage trait value of either Population 1 or 2, would result in a component of lineage sorting independent of the reference population. Similarly, the within-lineage trait change component (second term in eqn (1) and eqn (7)) can be made independent of the reference by using the average relative abundance of the lineages of the two populations. This results in a spatial version of the Price equation that is independent of the reference chosen, and partitions observed trait divergence between two spatially separated populations as:

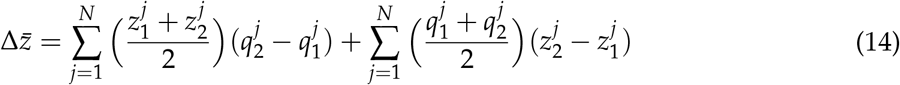

The first term on the right hand side of eqn (14) quantifies the observed trait divergence due to differences in relative abundances of the lineages between the two sites, while the second term quantifies observed trait divergence due to differences in trait values within lineages.

##### The reaction norm approach

As opposed to the spatial modification for the Price equation, we did not find a way to combine the plasticity, the constitutive evolution and evolution of plasticity components assessed from both directions (given in eqn (2) and (10)) to construct a spatial version for the reaction norm xapproach. The formula for evolution of plasticity is the same in both directions, and thus averaging would result in the same formula. However, taking an average plasticity and constitutive evolution component results in the same equation, implying that this metric would always lead to the conclusion that the contribution of plasticity and of genetic trait differentiation are equal, and thus becomes meaningless.

An alternative metric using reaction norms is provided by Ellner et al. (2011). While originally Ellner et al. (2011) quantified the ecological (impact of an environmental factor) and evolutionary (impact of the genetic component of a trait) to the change in an ecological response variable (e.g. population growth), this metric can also be used to partition the shift in a phenotypic trait *z* into main effects of evolution and ecology, i.e.

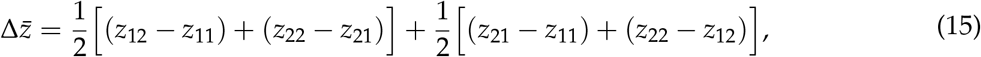

Similar as in the reaction norm approach, *z*_*kl*_ represents the trait value of the population at time *t*_*k*_ or site *k* in environmental condition *l*. A key difference between eqn (15) and the reaction norm approach presented by Govaert et al. (2016) is that the approach presented in Ellner et al. (2011) does not differentiate evolution into a component of constitutive evolution and a component of evolution of plasticity. The interpretation of the components of both approaches is thus different. The first term on the right hand side of eqn (15) sums the trait change in Population 1 (first part of the first term) and Population 2 (second part of the first term) due to a change in environmental condition from 1 to 2 divided by two, and can be seen as an average plasticity effect. The second term in eqn (15) sums the trait change in environment 1 (first part of the second term) and environment 2 (second part of the second term) between both populations divided by two, and can be seen as an average effect of genetic trait differentiation. This metric thus only estimates average effects of genetic trait differentiation and environmental (plasticity) and does not capture the full spectrum of evolutionary change (cf. evolution of plasticity is likely half included in each of the evolutionary and environmental components; Ellner et al. 2011). However, the advantage of this equation is that the contributions of evolution and ecology are the same (i.e. equal magnitude, but opposite in sign), and thus independent from whether one partitions trait shift from Population 1 to Population 2 or vice versa. This metric might thus be very suitable to quantify evolutionary and non-evolutionary contributions to spatially (undirected) trait shifts, at least if quantification of evolution of plasticity is not important.

### Application to empirical examples

Some spatial study systems do imply a directed trait change among spatially separated populations, such as studies investigating invasion history (e.g. stickleback colonisation of freshwater lakes; Bell et al. 2004; Le Rouzic et al. 2011), looking at range expansions where the peripheral population can be traced back to more central populations (e.g. Safriel et al. 1994; Volis et al. 2001; Swaegers et al. 2014) or using space-for-time substitutions to evaluate for example climate change (i.e. studies that infer temporal trends from populations that differ in age or in some temporally associated sequence; e.g. Etterson and Shaw 2001; Blois et al. 2013). We subsequently focus on studies that do not imply such a direction (but see Appendix E for an application of eco-evolutionary partitioning metrics to a study by Etterson and Shaw 2001). We applied eco-evolutionary partitioning metrics to three studies, illustrating which eco-evolutionary questions the modified metrics can address. In a first application to evolving meta-populations of *Ambystoma maculatum*, we determine whether contributions of plasticity and evolution depend on the selection pressure experienced (data obtained from Urban 2008). In a second application, we determine whether *Daphnia* species vary in their evolutionary and non-evolutionary responses to experimental environmental variation (data obtained from Weider et al. 2008). Last, we compared eco-evolutionary responses in two studies that evaluated a similar selection pressure, the addition of a predator species, to *Daphnia magna* in an experimental and a natural setting, to assess whether trait divergence in time and space can be structured by the same ecological and evolutionary processes (data obtained from De Meester 1996 and Cousyn et al. 2001).

#### Example 1: Do relative contributions of plasticity and evolution in a metapopulation of Ambystoma maculatum vary depending on the selection pressure experienced?

We evaluated whether *A. maculatum* populations originating from habitats with or without a predator (*A. opacum*) differed in the relative importance of plasticity and genetic trait differentiation for their deviation from the overall population mean larval size. Other possible research questions in a metapopulation could include, for example, how repeatable the contributions of plasticity and evolution to trait shift are when comparing an average predator-free *A. maculatum* population to each individual *A. maculatum* population co-existing with a predator or alternatively all pairs of predator-free and predator-present populations could be compared. These questions (addressed in Appendix E) would use different populations as reference points than the example presented here. We used the reaction norm equation from Approach 1 on a metapopulation of 18 *A. maculatum* populations originating from habitats varying in predation pressure (i.e. absence or presence of the predator *A. opacum*) collected by Urban (2008). Measurements of larval body mass of the 18 *A. maculatum* populations were assessed in a control and predator kairomone condition, which allows the construction of reaction norms (Fig. 4A). Using the reaction norm approach, we calculated contributions of plasticity, genetic trait differentiation and genetic differentiation in plasticity to shifts in prey body mass as measured in the laboratory for all 18 populations. Figure 4B and 4C show two alternative ways of visualising these results. If a division in groups of populations (here: those with and without predators) can be made, box-plots can be used to visualise differences in the relative contributions of the components between the two groups (Fig. 4B). However, one can also visualise the results in a triangle plot, which shows the actual values of the relative contribution of the three components for all 18 populations (Fig. 4C). Using different symbols for populations that originate from habitats with and without predator illustrates how these two groups differ in contributions of plasticity, genetic trait differentiation and genetic differentiation in plasticity to their trait deviation from the mean. Overall, plasticity contributed most to the observed deviations in mean prey body mass across the metapopulation, while genetic differentiation in plasticity had the smallest relative contribution. After categorising the 18 populations by whether or not they came from habitats with or without predation, we found that genetic differentiation in prey body mass varied significantly more in the populations where the predator was present compared to populations without predators (Levene’s test: *F*_1,16_ = 10.06, *p* = 0.006; Fig. 4B), and that the relative contribution of genetic differentiation in plasticity to body size was significantly larger in larvae originating from sites with predators compared to larvae from sites without predators (*t*-test: *t*_16_ = −2.21, *p* = 0.042; Fig. 4B). The narrow range in the relative contribution of genetic trait differentiation in prey body mass of the predator-free populations is depicted in the triangle plot by the grey zone (Fig. 4C).

**Figure 4:**
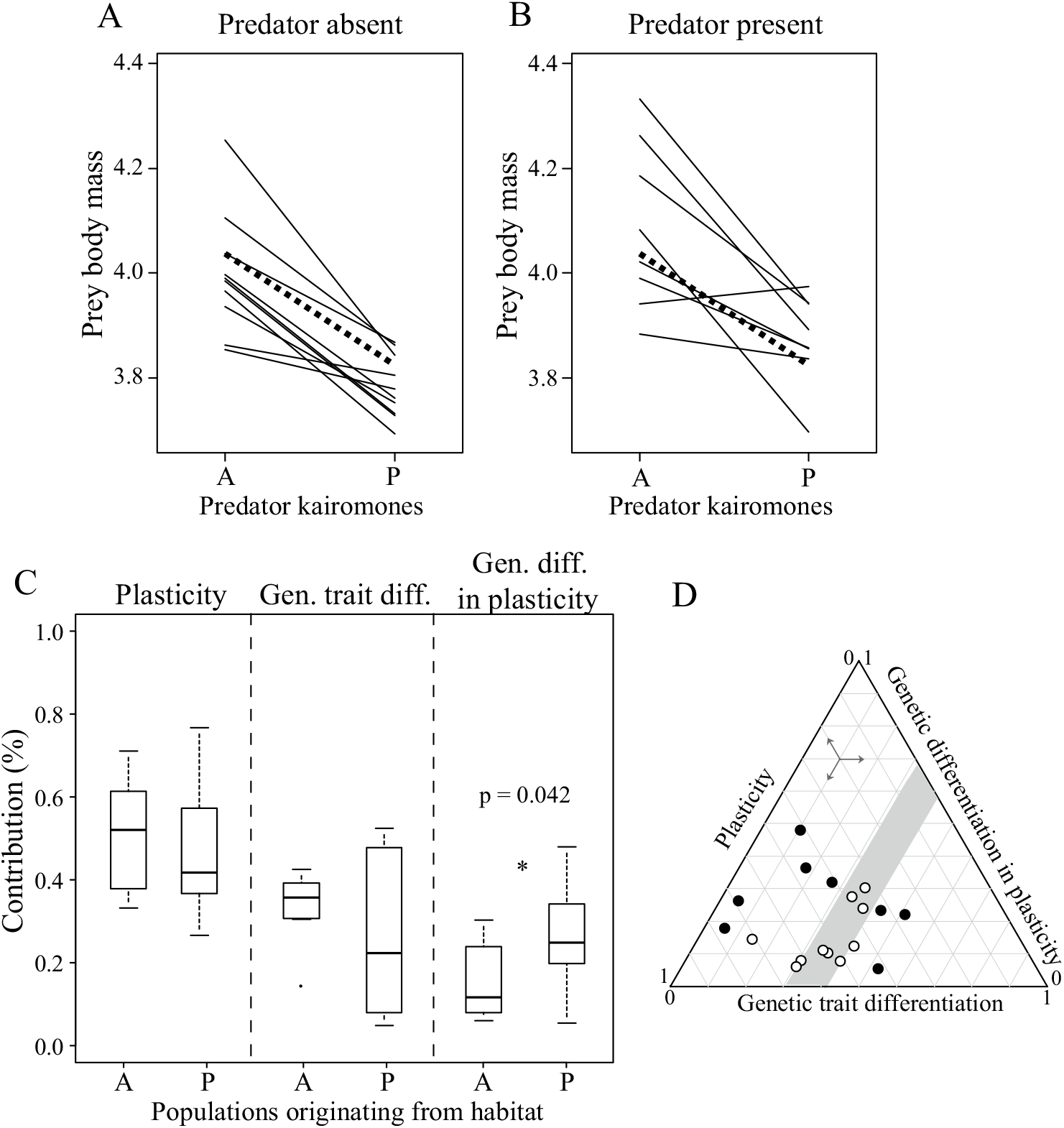
(A,B) Reaction norms for larval body mass of 18 populations of *A. maculatum* originating from sites where the predator *A. opacum* is either present or absent. Body mass of the larvae were measured in the absence (A) and presence (P) of predator kairomones. The dashed black line gives the group mean for all 18 populations. (C) Boxplots of the relative contributions of plasticity, genetic trait differentiation and genetic differentiation in plasticity relative to deviation in mean larval body mass using the reaction norm approach as described in Approach 1. Results are presented separately for the populations originating from habitats without (A) and with (P) the predator *A. opacum*. (D) Triangle plot showing the relative contributions of plasticity, genetic trait differentiation and genetic differentiation in plasticity. Filled (resp. unfilled) circles represent the *A. maculatum* populations originating from sites where the predator *A. opacum* is present (resp. absent). Data were extracted from Figure 3 in Urban (2008). Arrows on the triangle plot indicate how to read the coordinates of the points on the graph.

#### Example 2: Do relative contributions of clonal sorting and phenotypic plasticity vary among species and treatments?

We used the study by Weider et al. (2008) to quantify among-species variation in non-evolutionary and evolutionary contributions to shifts in age at first reproduction in an experiment involving three species of *Daphnia* cultured in a full-factorial design of high and low food quality and quantity. This example illustrates that spatially modified partitioning metrics can also be used to compare experimental treatments. In this example, we choose to construct a group mean (as opposed to pairwise comparisons among treatments) as this allows to associate non-evolutionary and evolutionary contributions to a single treatment effect. In an earlier study (Govaert et al. 2016), the same experimental data was used to determine the role of clonal and species sorting to temporal trait change in age at first reproduction in each experimental food condition using the Price-Reaction-Norm equation. While we would here use the Price-Reaction-Norm equation to assess non-evolutionary and evolutionary contributions to deviations in age at first reproduction, we cannot assess genetic trait differentiation and genetic differentiation in plasticity for this data set due to the experimental set-up. Hence, we decided to use a Price equation with interaction component to assess contributions of lineage sorting, within-lineage trait differentiation (here due to phenotypic plasticity) and their interaction to deviations in age at first reproduction for three *Daphnia* species (*D. pulex*, *D. pulicaria* and their hybrid) for four food conditions (LL: low-quality low-quantity; HL: high-quality low-quantity; LH: low-quality high-quantity; HH: high-quality high-quantity) at day 30 of a 90-day microcosm experiment. By plotting the absolute deviation from the average for the three components across treatments and species, we were able to detect whether species vary in the relative importance of these processes to the observed response to the experimental treatments and whether this variation among species differed among food conditions (Fig. 5). Overall, we found that within-lineage trait differentiation was the larger contributor (Fig. 5). Contributions of lineage sorting, within-lineage trait differentiation and their interaction were in opposite direction between the LL and HH treatments for all species except *D. pulicaria*. *D. pulex* and the hybrid species showed similar direction of the contributions for lineage sorting and within-lineage trait differentiation, where lower food quality was in opposite direction to high food quality conditions. By applying partitioning metrics to different species, we found that species varied in how the components were associated among treatments, indicating that different species may use different processes to respond to an environmental condition.

**Figure 5:**
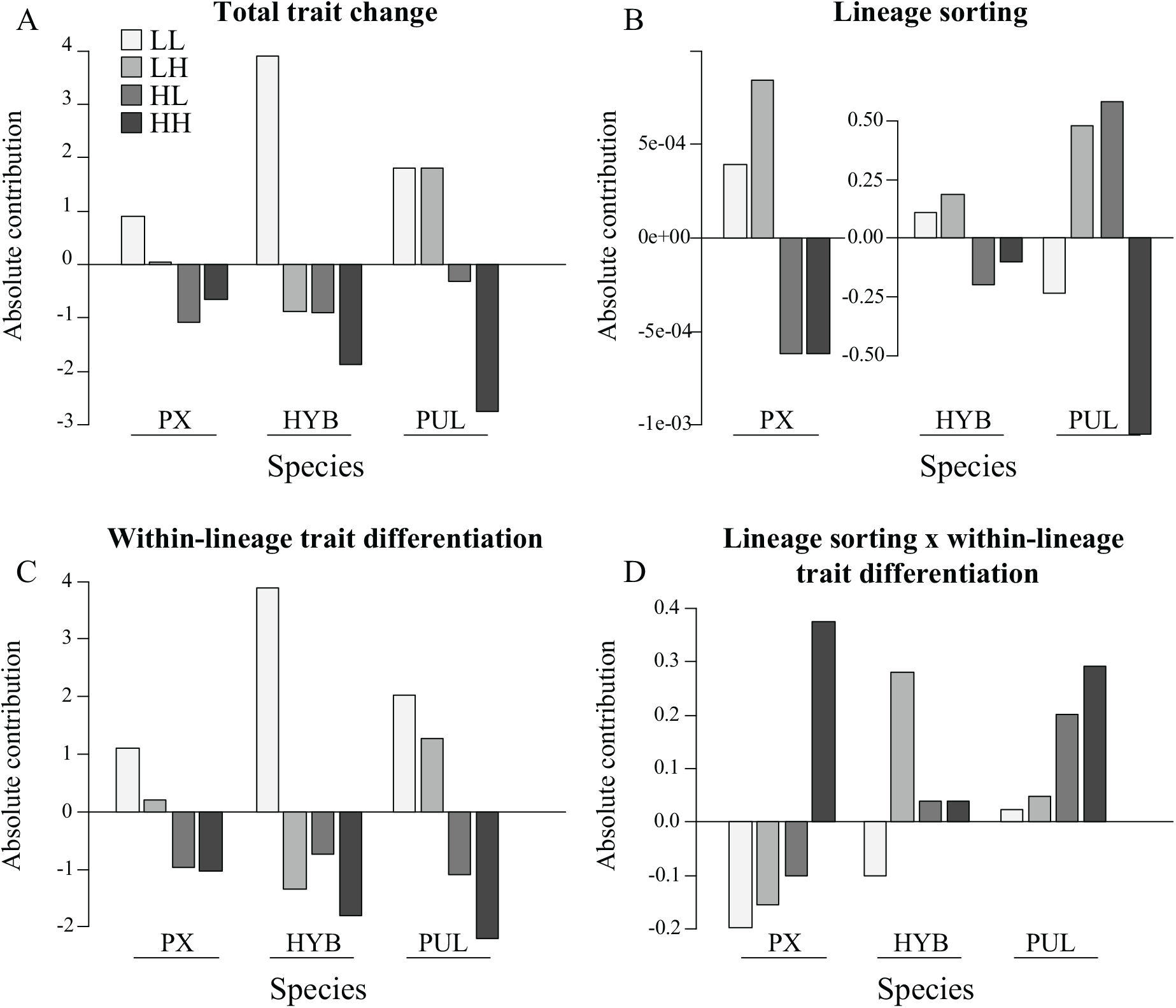
Graphical visualisation of (A) the total trait deviation from the group mean for each species (PX - *Daphnia pulex*, PUL - *Daphnia Pulicaria* and HYB - their hybrid) and treatment (low-quality low-quantity (LL), low-quality high-quantity (LH), high-quality low-quantity (HL), and high-quality high-quantity (HH)) and the absolute contributions of (B) shifts in lineage composition, (C) trait differentiation within lineages and (D) their interaction using the reaction norm approach.

#### Example 3: Are spatial and temporal trait divergence structured by the same processes? - A case study in Daphnia

De Meester (1996) and Cousyn et al. (2001) measured phenotypic responses in phototactic behaviour to different fish predation pressures (i.e. none, low and high) for three *Daphnia magna* (sub)populations in space and time (resurrection ecology approach), respectively. This provides an opportunity to investigate the similarity with respect to the importance of eco-evolutionary components in time and space. De Meester (1996) measured *D. magna* populations inhabiting different spatial locations with differing fish predation pressure (none, low, and high), while Cousyn et al. (2001) considered a single population that experienced temporal shifts in fish predation pressure (from none early 1970s to high fish predation pressure late 1970s and low mid 1980s). Both studies measured trait values for phototactic behaviour using a common garden experiment, exposing *D. magna* individuals of different clones from each (sub)population to a fish kairomone (mimicking high fish predation pressure) and a control (absence of fish predation threat) treatment. From these experiments, reaction norms could be constructed for each (sub)population (Fig. 6A-B), allowing the use of the reaction norm approach. These data sets have been previously used to illustrate the similarity of responses in phototactic behaviour in time and space by Freeman and Herron (2007), and we here explicitly quantify the importance of non-evolutionary and evolutionary processes in both the spatial and temporal setting. To facilitate this comparison, it was important to apply the same modification of the reaction norm approach for both data sets. In other words, although the resurrection ecology reconstruction of Cousyn et al. (2001) allows application of a directional partitioning metric (see Stoks et al. 2016), we chose to apply an undirected metric to enable direct comparison of the magnitudes of the components with the spatial study. We used the reaction norm approach as described in Approach 1 to calculate deviations of populations from the overall group mean (e.g. the mean across all individuals from all populations within a study). By comparing the (sub)populations to this group mean, we could assess if (sub)populations experiencing a similar predation pressure also similarly deviated from the group mean in their non-evolutionary and evolutionary contributions. To test the latter, we performed a bootstrap analysis resampling the data with replacement, and recalculating the contributions of plasticity, genetic trait differentiation and genetic differentiation in plasticity. We then compared the 95% confidence intervals of the relative (Fig. 6C) and absolute (Fig. 6D-L) contributions of plasticity, genetic trait differentiation and genetic differentiation in plasticity between pairs of (sub)populations that experienced similar fish predation pressure. We found that all 95% confidence intervals obtained from the bootstrapping overlapped for the absolute contributions of plasticity, genetic trait differentiation and genetic differentiation in plasticity, indicating a similar range of the absolute contributions of these processes between pairs of (sub)populations that experienced similar fish predation pressure. We thus found that a shared selection pressure resulted in a similar allocation of trait change across the ecological, evolutionary, and eco-evolutionary contributions and this result was independent on the approach used (detailed in Appendix F). Our analysis thus suggests that in this case adaptation through time and across space is achieved through similar combinations of mechanisms. This also suggests that the spatial differentiation observed in De Meester (1996) could in principle be achieved in a time span of a few years (the time span of the resurrection ecology study).

**Figure 6:**
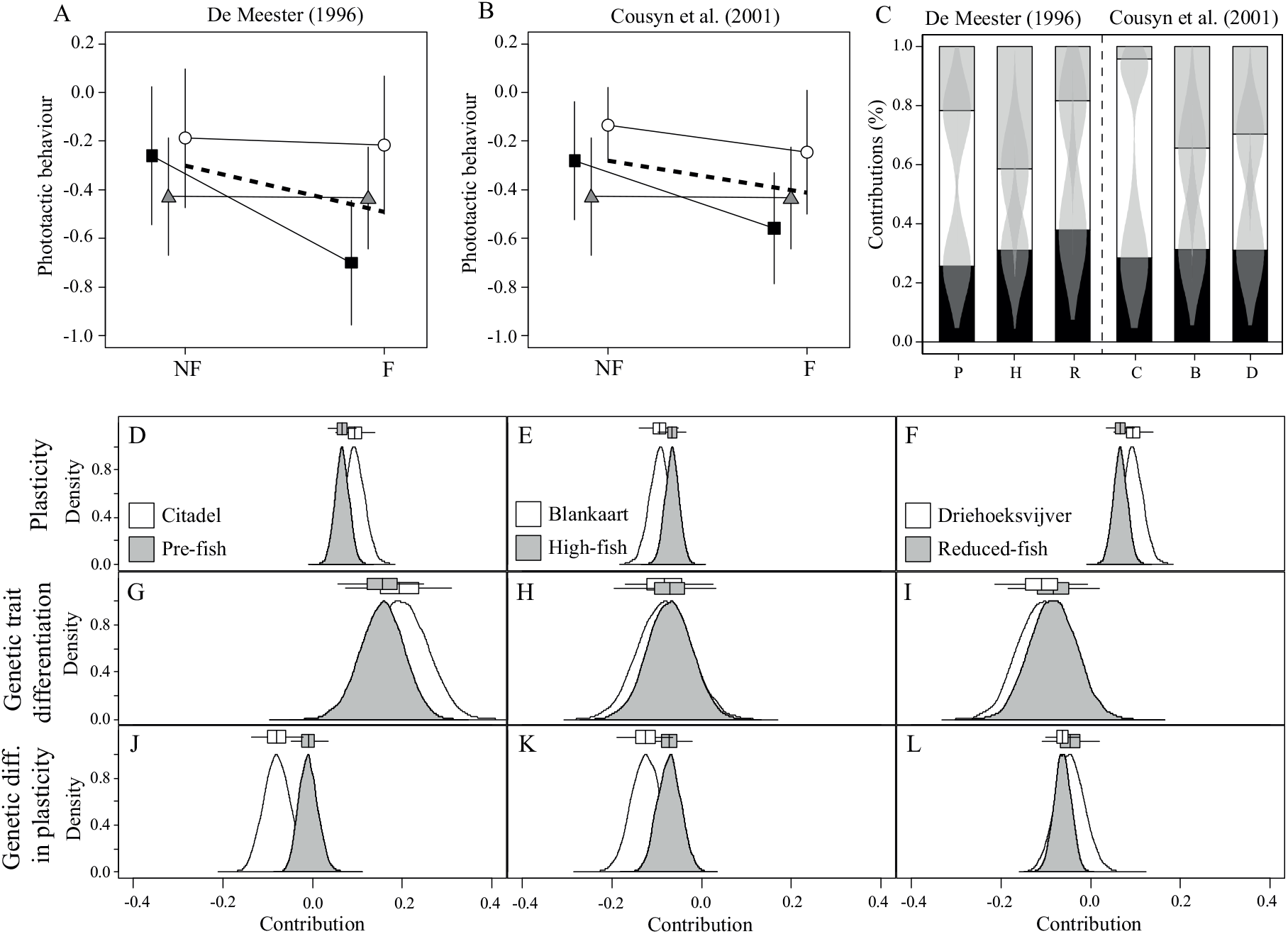
Reaction norms of phototactic behaviour of three *D. magna* populations (A) originating from three different ponds corresponding to no (Citadelpark, C; circles), high (Blankaart, B; squares) and low fish predation pressure (Driehoeksvijver, D; triangles) and (B) from the same pond Oud-Heverlee Zuid at three different time periods corresponding to no (Pre-fish; circles), high (High-fish; squares) and reduced (Reduced-fish; triangles) fish predation pressure measured in a control (NF) and fish kairomone (F) condition (data obtained from Figure 1 in De Meester 1996 and from Figure 2 in Cousyn et al. 2001). (C) Relative contributions of plasticity (black), genetic trait differentiation (white) and genetic differentiation in plasticity (grey) and their density distribution obtained from bootstrap analysis given as a violin plot within the bars (shaded grey area) to observed difference in phototactic behaviour from the group mean to each population from De Meester (1996) - (population C, B, or D) - and from Cousyn et al. (2001) - Pre-fish (P), High-fish (H) or Reduced-fish (R) - using the reaction norm approach as explained in Approach 1. (D-L) Bootstrap distributions of the absolute contributions of plasticity (D-F), genetic trait differentiation (G-I) and genetic differentiation in plasticity (J-L) for the populations from De Meester (1996) (white), and from Cousyn et al. (2001) (grey).

## Discussion

The phenotypic distribution of spatially separated populations and communities can be structured by non-evolutionary (phenotypic plasticity at the individual level; species sorting at the community level) and evolutionary (genetic differentiation) processes (Via and Lande 1985; Kawecki and Ebert 2004; Fox and Harder 2015; Govaert et al. 2016; Logan et al. 2016). However, evolutionary and non-evolutionary contributions to trait divergence among populations or communities separated in space cannot always be quantified using the same methods as for population or community trait change in time. Temporal studies may ask how evolutionary and non-evolutionary processes combine to structure trait shifts from one time point to the next, while a spatial study instead seeks to quantify evolutionary and non-evolutionary contributions to among-site trait divergence. While spatial studies cannot reconstruct how traits changed through time or assess changes in community composition resulting from species extinctions and colonizations mediated through evolution (a temporal process), they can quantify to what extent plasticity, genetic trait differentiation and species sorting combine to explain among-site differences in trait values. In this study, we illustrated three ways to adjust the Price equation (Price 1970; Price 1972), the reaction norm approach (Ellner et al. 2011; Govaert et al. 2016) and the Price-Reaction-Norm equation (Govaert et al. 2016) to match spatial, undirected comparisons of trait shifts. We then applied some of the metrics to published case studies of empirical data to illustrate the diverse array of questions that can be addressed when the metrics are matched to the question of interest.

We presented three approaches to quantify ecological and evolutionary contributions to spatial trait divergence using existing partitioning metrics. The first approach involves constructing a group mean that represents a known ancestral state or control treatment, and evaluating how individual sub-populations differ from that group average. The choice of the group mean depends on the researcher’s question – it could represent a metapopulation average, or a global mean across all levels of an experiment. Once the partition analysis is conducted and eco-evolutionary contributions to trait shifts from the group average are calculated for each sub-population, these contributions can be compared among sub-populations, but can also be linked to population-specific characteristics. For example, among-population genetic differentiation is expected when sub-populations are spatially isolated (Wright 1943; Bohonak 1999), and the degree of evolutionary trait shift can be compared with the population’s degree of spatial isolation. More specific eco-evolutionary hypotheses, for example whether or not populations with a longer history of exposure to a selection pressure (Ghalambor et al. 2007; Oostra et al. 2018) or different selection pressures (Huang and Agrawal 2016), or with a different degree of environmental variability (Reed et al. 2010; Chevin and Hoffmann 2017) demonstrate a larger fraction of genetic evolution versus plasticity could be tested by comparing the different fractions to these site properties. The second and third approach we presented do not require designating an ancestral or control reference. They instead use different methods of averaging and consider comparisons between pairs of populations. The second approach calculates the ecological and evolutionary components of among-population trait differences by treating each population in the pair as the reference, then averaging the two resulting absolute values for each component. It thus produces an overall assessment of how the importance of evolutionary and non-evolutionary processes varies among a set of spatially separated populations or communities. The third approach also does not require choosing one population as the initial state. Depending on the metric used, it instead averages either the two population’s trait values and/or abundance values when calculating eco-evolutionary components. Because in the second approach, the averaging is across absolute values of the resulting ecological and evolutionary contributions, the resulting eco-evolutionary fractions do not sum to the single observed difference in traits between the two populations.

These spatial eco-evolutionary partitioning metrics were applied to data from three existing studies to illustrate how they can answer eco-evolutionary research questions. Spatial partitioning metrics can identify spatial structure in the relative contributions of evolutionary and non-evolutionary processes to trait variation in natural landscapes. Spatial variation in abiotic conditions and ecological interactions may result in spatially divergent selection strengths, producing distinct evolutionary trajectories among populations (and between coevolving species, i.e. the geographic mosaic of coevolution, Thompson 1999; Thompson 2005). These different selection pressures in a heterogeneous landscape might result in varying contributions of non-evolutionary and evolutionary processes, and these contributions could further depend on the population or community identity, or on the focal species studied. The case study of Urban (2008) indicated that populations of *A. maculatum* living in the presence of the predator *A. opacum* had substantially higher contributions of genetic differentiation in plasticity and showed larger variation in their genetic trait differentiation than populations living in the absence of the predator. The populations living in the absence of the predator showed a strikingly narrow range in the relative contribution of genetic trait differentiation in larval body size. Similarly, the case study of Weider et al. (2008) indicated that eco-evolutionary contributions differed depending on experimental treatments, but also among species, although there is currently no clear expectations for when this dependence is more or less likely. Nevertheless, the indication of context-dependent eco-evolutionary processes is important for future research in the field of eco-evolutionary dynamics.

It currently is unknown to what extent the magnitude of eco-evolutionary contributions to population and community trait change are predictable and repeatable. In this study we used spatial eco-evolutionary partitioning metrics to compare how *D. magna* traits responded to the presence of fish predators, and found a similarity in eco-evolutionary responses in phototatic behaviour between populations with spatial variation in this selection pressure and populations with temporal variation. Although the similarity in spatial and temporal trait shifts has previously been explored for the *D. magna* populations used in this study (Freeman and Herron 2007), our application explores whether eco-evolutionary contributions to spatial and temporal trait shifts are also repeatable in space and time. This is an intriguing possibility, because it has previously been hypothesized that trait evolution dynamics in space and time can be similar (e.g. Frank 1991; Gandon et al. 2008). Our results indicate this similarity may also be reflected when considering both plastic and genetic trait responses to selection pressures. We anticipate further comparative studies are needed to establish whether this is a repeatable pattern.

Directionality is not always lacking in spatial research studies. Studies that use space-for-time substitutions to evaluate how traits shift from cooler to warmer climates (Etterson and Shaw 2001; Dinh Van et al. 2014; Janssens et al. 2014) and studies that compare trait shifts from an ancestral to an invasive populations are some examples where this directionality of the trait shift is clear (Bell et al. 2004; Le Rouzic et al. 2011). These studies can thus use existing partitioning metrics (that assign an ancestral and dependent state) to understand how eco-evolutionary processes contribute to these trait shifts. Integrating the assessment of contributions of genetic and non-genetic variation to the expected trait change can potentially inform future conservation planning (Moritz 2002). For instance, if the expected trait change is mainly due to plasticity and increases the organism’s fitness, it is more likely that the population will experience a short-term positive response to the changing environment. This provides time for the population to genetically respond to the changing environment, which might eventually be essential for its future persistence (Bradshaw and Holzapfel 2006; Gienapp et al. 2008). However, if the expected trait change is mainly due to evolution, conservation management can take the amount of genetic variation and evolutionary potential in the target populations into consideration. Conservation of evolutionary potential means taking measures to maintain or increase genetic diversity (such as habitat restoration or creation, supporting a well-connected metapopulation), while taking the risk of outbreeding depression into account (Fenster and Dudash 1994). It is important to realize that the genetic structure of the population changes as it adapts to the environmental change, and this might impact its future response to other stressors (e.g. in the case of genetic erosion; Harlan 1975; Bijlsma and Loeschcke 2012). Conservation efforts that focus only on species diversity may potentially overlook the importance of intraspecific and genetic trait variation and how this can influence ecological dynamics (Lande 1988; Moritz 1994; Hughes et al. 1997; Frankham et al. 2002; Moritz 2002; Palkovacs et al. 2012; Des Roches et al. 2018).

Intraspecific trait variation might be critical for population dynamics, community structure and ecosystem functioning in a wide range of settings (Mimura et al. 2017; Des Roches et al. 2018). However, the importance of intraspecific variation is likely to vary. Some important questions that remain to be answered next are: how much of this intraspecific trait variation is due to genetic trait variation?, and to what extent does the ecological importance of genetic trait variation differ across landscapes properties such as connectivity, across biotic and abiotic environmental gradients, and in response to interactions with other species? The approaches outlined here to convert existing eco-evolutionary partitioning metrics to appropriately accommodate spatially structured (undirected) trait data will facilitate future attempts to determine associations between among-site variation in evolutionary and non-evolutionary components and properties of the landscape, the environment or the study species. Numerous studies compare trait distributions among communities (Cornwell and Ackerly 2009; Vellend 2016; Kenitz et al. 2018) or population genetic structure among populations (Marten et al. 2006; Gomez-Uchida et al. 2009; Short and Caterino 2009; Olsen et al. 2011; Ackerman et al. 2013). However, there are very few studies that collected the necessary data to decompose all potential sources of trait shifts at the community level. We anticipate an increase in the number of studies that attempt to combine surveys of genetic and non-genetic trait variation at the population level with species composition and associated trait shifts at the community level. The data gathered by such studies can be used to quantify the contributions of evolutionary and non-evolutionary processes to among-site variation in community trait values, quantifying the structure of the evolving metacommunity. We therefore predict an increasing scope for the application of the metrics proposed in this study.

## Acknowledgments

This research was supported by KU Leuven Research Fund projects PF/2010/07 and C16/17/002, Belspo IUAP project SPEEDY P7/04, FWO projects G0B9818 and G0C3818, KU Leuven Research Fund F+ 13036 fellowship awarded to JHP, and an IWT PhD fellowship awarded to LG. LG was also supported by the University of Zurich Research Priority Program on ‘Global Change and Biodiversity’.

