## Supplementary Information for "Quantifying eco-evolutionary contributions to trait divergence in spatially structured systems"

Lynn Govaert[1,2,3*]

Jelena H. Pantel[1,4]

Luc De Meester[1]

1. Laboratory of Aquatic Ecology, Evolution and Conservation, University of Leuven, Ch. Deberiotstraat 32, B-3000 Leuven, Belgium;

2. Department of Evolutionary Biology and Environmental Studies, University of Zurich, Winterthurerstrasse 190, CH-8057 Zürich, Switzerland;

3. Swiss Federal Institute of Aquatic Science and Technology, Department of Aquatic Ecology, Überlandstrasse 133, CH-8600 Dübendorf, Switzerland;

4. Department of Computer Science, Mathematics, and Environmental Science, The American University of Paris, 6 rue du Colonel Combes, 75007 Paris, France

*Manuscript elements*: Appendices A-F including Figure C1, D1 and F1-F8

*Keywords*: eco-evolutionary dynamics, partitioning metrics, spatial trait variation, population, trait change

### Appendix A: Extension to spatially separated communities

While our focus is toward populations, each of the discussed approaches in the main text can be extended to the community level as there exists a community version of the Price equation, reaction norm approach and Price-Reaction-Norm equation. We advise the reader to first read through the main text before going through this appendix.

We start with briefly showcasing the extension of the spatially modifications to partition ecological and evolutionary contributions to trait divergence among a set of $m$ spatially separated communities, uniquely indexed by $k \in \{1, \ldots, m\}$. However, we first briefly give an overview on the Price equation (Price 1970), the reaction norm approach (Stoks et al. 2016; Govaert et al. 2016) and the Price-Reaction-Norm equation (Govaert et al. 2016) at the community level used in a temporal context (i.e. community trait change occurs between consecutive time points). Similarly as at the population level, these require different set of data information, and hence they differ in the components they can assess.

The Price equation can be used to partition community trait change between two time points of a community consisting of $s$ asexually reproducing species, uniquely indexed by $i \in \{1, \ldots, s\}$, where each species $i$ consists of $N_i$ genetic lineages, uniquely indexed by $j \in \{1, \ldots, N_i\}$ into species sorting, lineage sorting and trait change within lineages, i.e.

$$\Delta \bar{z} = \sum_{i=1}^{s} z_2^i (q_2^i - q_1^i) + \sum_{i=1}^{s} q_1^i \sum_{j=1}^{N_i} z_2^{ij} \Big( \frac{q_2^{ij}}{q_2^i} - \frac{q_1^{ij}}{q_1^i} \Big) + \sum_{i=1}^{s} q_1^i \sum_{j=1}^{N_i} \frac{q_1^{ij}}{q_1^i} (z_2^{ij} - z_1^{ij}), \tag{A1}$$

where $z_k^{ij}$ (resp. $q_k^{ij}$) is the average trait value (resp. relative abundance within the community) of genetic lineage $j$ of species $i$ at time point $t_k$, and $z_k^i$ (resp. $q_k^i$) is the average trait value (resp. relative abundance) of species $i$.

For the reaction norm approach we assume that the community consists of $s$ species, uniquely indexed by $i \in \{1, \ldots, s\}$ inhabiting two distinct environments (e.g. low or high temperature, absence or presence predator) at the two time points of measurement. Furthermore, information on the average trait value of each species of the community at both time points at the two environmental conditions should be available. This allows construction of species mean reaction norms. The reaction norm approach can then be used to partition community trait change into the following components:

$$\begin{aligned}\Delta\bar{z} = &\sum_{i=1}^{s} q_1^i(z_{12}^i - z_{11}^i) + \sum_{i=1}^{s} q_1^i(z_{21}^i - z_{11}^i)\\ &+ \sum_{i=1}^{s} q_1^i\big([z_{22}^i - z_{21}^i] - [z_{12}^i - z_{11}^i]\big)\\ &+ \sum_{i=1}^{s} z_{12}^i(q_2^i - q_1^i) + \sum_{i=1}^{s}(q_2^i - q_1^i)(z_{21}^i - z_{11}^i)\\ &+ \sum_{i=1}^{s}(q_2^i - q_1^i)\big([z_{22}^i - z_{21}^i] - [z_{12}^i - z_{11}^i]\big),\end{aligned} \tag{A2}$$

where $z_{kl}^i$ is the $i$th species mean trait value of the community at time point $t_k$ in the environmental condition $l$. The first term on the right hand side of eqn (A2) reflects the plasticity component. The second term represents constitutive evolution, the third term evolution of plasticity, the fourth term species sorting, and the fifth and sixth term interactions between species sorting and constitutive evolution and between species sorting and evolution of plasticity.

Last, the Price-Reaction-Norm equation combines features of the Price equation and of the reaction norm approach (Govaert et al. 2016). Consider a community consisting of $s$ asexually reproducing species, uniquely indexed by $i \in \{1, \ldots, s\}$, where each species $i$ consists of $N_i$ genetic lineages, uniquely indexed by $j \in \{1, \ldots, N_i\}$ measured at two time points that can be linked to two distinct environments. For the Price-Reaction-Norm equation we require to have information on the average trait values and relative abundances of each genetic lineage $j$ of a species $i$ of both time points measured at the two environmental conditions. The Price-Reaction-Norm equation can then be used to disentangle community trait change into eight additive components, i.e.

$$
\begin{aligned}
\Delta\bar{z} = & \sum_{i=1}^{s} q_1^i \sum_{j=1}^{N_i} \frac{q_1^{ij}}{q_1^i}(z_{12}^{ij} - z_{11}^{ij}) + \sum_{i=1}^{s} q_1^i \sum_{j=1}^{N_i} z_{22}^{ij}\Big(\frac{q_2^{ij}}{q_2^i} - \frac{q_1^{ij}}{q_1^i}\Big) \\
& + \sum_{i=1}^{s} q_1^i \sum_{j=1}^{N_i} \frac{q_1^{ij}}{q_1^i}(z_{21}^{ij} - z_{11}^{ij}) + \sum_{i=1}^{s} q_1^i \sum_{j=1}^{N_i} \frac{q_1^{ij}}{q_1^i}([z_{22}^{ij} - z_{21}^{ij}] - [z_{12}^{ij} - z_{11}^{ij}]) \\
& + \sum_{i=1}^{s} z_{12}^i (q_2^i - q_1^i) + \sum_{i=1}^{s} (q_2^i - q_1^i) \sum_{j=1}^{N_i} z_{22}^{ij}\Big(\frac{q_2^{ij}}{q_2^i} - \frac{q_1^{ij}}{q_1^i}\Big) \\
& + \sum_{i=1}^{s} (q_2^i - q_1^i) \sum_{j=1}^{N_i} \frac{q_1^{ij}}{q_1^i}(z_{21}^{ij} - z_{11}^{ij}) \\
& + \sum_{i=1}^{s} (q_2^i - q_1^i) \sum_{j=1}^{N_i} \frac{q_1^{ij}}{q_1^i}([z_{22}^{ij} - z_{21}^{ij}] - [z_{12}^{ij} - z_{11}^{ij}]),
\end{aligned}
\qquad \text{(A3)}
$$

where $z_{kl}^{ij}$ (resp. $q_k^{ij}$) is the average trait value (resp. relative abundance within the community) of genetic lineage $j$ of species $i$ in the community at time point $t_k$ measured at environmental condition $l$, and $z_{kl}^i$ (resp. $q_k^i$) is the average trait value (resp. relative abundance) of species $i$ of the community at time point $t_k$ measured at environmental condition $l$. The terms on the right hand side of eqn (A3), in order of appearance, reflect phenotypic plasticity, lineage sorting, genetic trait change within lineages, evolution of plasticity within lineages, species sorting, species sorting $\times$ lineage sorting, species sorting $\times$ genetic trait change within lineages and species sorting $\times$ evolution of plasticity.

We next describe an extension of Approach 1, Approach 2 and 3 described in the main text to community trait data. For Approach 1, we should calculate a group mean across the $m$ spatially separated communities, and then use the temporal versions of the eco-evolutionary partitioning metrics to calculate deviations from this group mean to each community. For Approach 2, we calculate the non-evolutionary and evolutionary contributions to trait shifts among pairs of communities, for example, from Community 1 to Community 2, and then from Community 2 to Community 1 using either the Price equation, the reaction norm approach or the Price-Reaction-Norm equation. The absolute values of these contributions are then averaged to give the overall relative importance of ecological and evolutionary processes. Approach 3 involves a mathematical modification of the metrics to become independent of a reference that is conceptually similar to the modifications for each metric previously shown at the population level.

#### *Approach 1: Deviation from group mean*

In the main text we showed how a group mean can be created for the Price equation, the reaction norm approach and the Price-Reaction-Norm equation at the population level. This group mean can then be used to calculate non-evolutionary and evolutionary contributions to trait deviations for each population. Similarly, such a group mean can be constructed for a set of $m$ spatially separated communities.

##### *The Price equation*

For the Price equation, a group mean for $m$ spatially separated communities can be constructed by first calculating the average lineage trait values $z_a^{ij}$ and relative abundances $q_a^{ij}$ for each lineage $j$ of species $i$, similarly as in eqn (3) in main text. Using these average trait values and abundances, one can calculate a species mean trait value ($z_a^i$) and relative abundance ($q_a^i$) for each species $i$:

$$z_a^i = \sum_{j=1}^{N_i} \frac{q_a^{ij}}{q_a^i} z_a^{ij} \text{ and } q_a^i = \sum_{j=1}^{N_i} q_a^{ij}.$$

The community version of the Price equation (eqn (A1)) can then be used to quantify the contribution of species sorting, lineage sorting and trait deviation within lineages to observed community trait deviation from the group mean to each Community $k$ where $k \in \{1, \ldots, m\}$:

$$\Delta \bar{z}_{C_a \to C_k} = \sum_{i=1}^{s} z_k^i (q_k^i - q_a^i) + \sum_{i=1}^{s} q_a^i \sum_{j=1}^{N_i} z_k^{ij} \left( \frac{q_k^{ij}}{q_k^i} - \frac{q_a^{ij}}{q_a^i} \right) + \sum_{i=1}^{s} q_a^i \sum_{j=1}^{N_i} \frac{q_a^{ij}}{q_a^i} (z_k^{ij} - z_a^{ij}), \tag{A4}$$

The first term on the right hand side of eqn (A4) quantifies the community trait deviation due to species sorting, the second term quantifies the community trait deviation due to lineage sorting within species, and the last term quantifies the community trait deviation caused by within-lineage trait deviation.

*The reaction norm approach*

For the reaction norm approach a group mean can be constructed by calculating $z^i_{al}$ (i.e. the average trait value of species $i$ at environmental condition $l \in \{1, 2\}$ across the total set of $m$ communities), $z^i_{ka}$ (i.e. the average trait value of species $i$ from a specific Community $k$ at the average environmental condition $a$), $z^i_{aa}$ (i.e. the average trait value of species $i$ at the average environmental condition $a$), and $q^i_a$ (i.e. the average relative abundance of species $i$ across the total set of $m$ communities) as follows:

$$z^i_{al} = \frac{1}{m}\sum_{k=1}^{m} z^i_{kl}, \quad z^i_{ka} = \frac{z^i_{k1} + z^i_{k2}}{2}, \quad z^i_{aa} = \frac{1}{m}\sum_{k=1}^{m} z^i_{ka} \quad \text{and} \quad q^i_a = \frac{1}{m}\sum_{k=1}^{m} q^i_k \tag{A5}$$

The reaction norm approach as given in eqn (A2) can then be used to decompose each community trait's deviation from the group mean as follows:

$$\begin{aligned}\Delta\bar{z}_{C_a \to C_k} = &\sum_{i=1}^{s} q^i_a(z^i_{ak} - z^i_{aa}) + \sum_{i=1}^{s} q^i_a(z^i_{ka} - z^i_{aa}) \\ &+ \sum_{i=1}^{s} q^i_a\big([z^i_{kk} - z^i_{ka}] - [z^i_{ak} - z^i_{aa}]\big) + \sum_{i=1}^{s} z^i_{ak}(q^i_k - q^i_a) \\ &+ \sum_{i=1}^{s}(q^i_k - q^i_a)(z^i_{ka} - z^i_{aa}) + \sum_{i=1}^{s}(q^i_k - q^i_a)\big([z^i_{kk} - z^i_{ka}] - [z^i_{ak} - z^i_{aa}]\big),\end{aligned} \tag{A6}$$

with $l \in \{1, 2\}$ and $k \in \{1, \ldots, m\}$. The first term on the right hand side of eqn (A6) is the plasticity component. The second term is the trait deviation due to genetic trait differentiation of the different species for each Community $k$. The third term reflects genetic differentiation in plasticity. The fourth term is the trait deviation due to species sorting. The fifth and sixth term are interactions between species sorting and genetic trait differentiation and between species sorting and genetic differentiation in plasticity.

*The Price-Reaction-Norm equation*

Last, for the Price-Reaction-Norm equation, the group mean is constructed by first calculating the average trait values and relative abundances for the genetic lineages of each species $i$ in each community $k$ (described in eqn (B2), Appendix B), i.e. $z^{ij}_{al}$ (the average trait value of lineage $j$ of species $i$ from the average community at environmental condition $l \in \{1,2\}$), $z^{ij}_{ka}$ (the average trait value of lineage $j$ of species $i$ from a specific Community $k$ at the average environmental condition $a$), $z^{ij}_{aa}$ (the average trait value of lineage $j$ of species $i$ from the average community at the average environmental condition $a$), and $q^{ij}_{a}$ (the average frequency of lineage $j$ of species $i$ among the communities) and this for each species $i$. From these average lineage trait values we can calculate the species trait values $z^{i}_{al}$, $z^{i}_{ka}$ and $z^{i}_{aa}$ and the species relative abundance $q^{i}_{a}$ as follows:

$$z^{i}_{al} = \sum_{j=1}^{N_i} \frac{q^{ij}_{a}}{q^{i}_{a}} z^{ij}_{al}, \qquad z^{i}_{ka} = \sum_{j=1}^{N_i} \frac{q^{ij}_{a}}{q^{i}_{a}} z^{ij}_{ka}, \qquad z^{i}_{aa} = \sum_{j=1}^{N_i} \frac{q^{ij}_{a}}{q^{i}_{a}} z^{ij}_{aa}, \qquad \text{and} \qquad q^{i}_{a} = \sum_{j=1}^{N_i} q^{ij}_{a}.$$

The PRN equation as given in eqn (A3) can then be used to decompose the observed trait deviation from the group mean to Community $k$ into the following components:

$$\begin{aligned}
\Delta\bar{z}_{C_a \to C_k} = & \sum_{i=1}^{s} q^{i}_{a} \sum_{j=1}^{N_i} \frac{q^{ij}_{a}}{q^{i}_{a}} (z^{ij}_{ak} - z^{ij}_{aa}) + \sum_{i=1}^{s} q^{i}_{a} \sum_{j=1}^{N_i} z^{ij}_{kk} \Big( \frac{q^{ij}_{k}}{q^{i}_{k}} - \frac{q^{ij}_{a}}{q^{i}_{a}} \Big) \\
& + \sum_{i=1}^{s} q^{i}_{a} \sum_{j=1}^{N_i} \frac{q^{ij}_{a}}{q^{i}_{a}} (z^{ij}_{ka} - z^{ij}_{aa}) + \sum_{i=1}^{s} q^{i}_{a} \sum_{j=1}^{N_i} \frac{q^{ij}_{a}}{q^{i}_{a}} \big( [z^{ij}_{kk} - z^{ij}_{ka}] - [z^{ij}_{ak} - z^{ij}_{aa}] \big) \\
& + \sum_{i=1}^{s} z^{i}_{ak} (q^{i}_{k} - q^{i}_{a}) + \sum_{i=1}^{s} (q^{i}_{k} - q^{i}_{a}) \sum_{j=1}^{N_i} z^{ij}_{kk} \Big( \frac{q^{ij}_{k}}{q^{i}_{k}} - \frac{q^{ij}_{a}}{q^{i}_{a}} \Big) \\
& + \sum_{i=1}^{s} (q^{i}_{k} - q^{i}_{a}) \sum_{j=1}^{N_i} \frac{q^{ij}_{a}}{q^{i}_{a}} (z^{ij}_{ka} - z^{ij}_{aa}) \\
& + \sum_{i=1}^{s} (q^{i}_{k} - q^{i}_{a}) \sum_{j=1}^{N_i} \frac{q^{ij}_{a}}{q^{i}_{a}} \big( [z^{ij}_{kk} - z^{ij}_{ka}] - [z^{ij}_{ak} - z^{ij}_{aa}] \big),
\end{aligned} \tag{A7}$$

with $l \in \{1,2\}$ for the two environmental conditions a community originates from and $k \in \{1,\ldots,m\}$. The first term on the right hand side of eqn (A7) is the trait deviation due to plasticity. The second, third and fourth term are evolutionary components and quantify the trait deviation due to lineage sorting, within-lineage genetic trait differentiation and genetic differentiation in plasticity within lineages from the group mean. The fifth term is the trait deviation due to species sorting, and the sixth, seventh and eighth term are interactions between species sorting and lineage sorting, between species sorting and genetic trait differentiation within lineages, and between species sorting and genetic differentiation in plasticity, respectively.

##### *Approach 2: Average of components in both directions of comparison*

In the second approach, detailed in the main text for spatially separated populations, we suggested to partition trait divergence between pairs of populations using each population in the pair as a reference. In the case of $m$ spatially separated communities, this would mean that ecological and evolutionary contributions would be quantified to the trait divergence between pairs of communities by taking either community in the pair as a reference (i.e. quantifying ecological and evolutionary contributions to community trait shift from Community $k$ to Community $r$ and then repeating the calculation taking Community $r$ as the reference). Absolute values of ecological and evolutionary components obtained from either the community version of the Price equation (eqn (A1)), reaction norm approach (eqn (A2)) or Price-Reaction-Norm equation (eqn (A3)) for each comparison could then be averaged to get an overall quantification of the importance of evolutionary and ecological processes to the observed trait divergence. We next illustrate how these components are calculated for the Price equation, reaction norm approach and Price-Reaction-Norm equation. For clarity, we use index 1 and 2 instead of $k$ and $r$ for indicating the two communities used in the pairwise comparison.

###### *Metric 1: The Price equation*

By using index 1 and 2, equation (A1), given previously, quantifies the evolutionary and non-evolutionary contributions to the observed community trait shift going from Community 1 to Community 2. Hence, using Community 1 as a reference. Using Community 2 as a reference, we can similarly use the community version of the Price equation to quantify evolutionary and non-evolutionary contributions the community trait divergence ($\Delta\bar{z}_{C_2\to C_1} = \bar{z}_1 - \bar{z}_2$) as follows:

$$\Delta\bar{z}_{C_2\to C_1} = \sum_{i=1}^{s} z_1^i(q_1^i - q_2^i) + \sum_{i=1}^{s} q_2^i \sum_{j=1}^{N_i} z_1^{ij}\Big(\frac{q_1^{ij}}{q_1^i} - \frac{q_2^{ij}}{q_2^i}\Big) + \sum_{i=1}^{s} q_2^i \sum_{j=1}^{N_i} \frac{q_2^{ij}}{q_2^i}(z_1^{ij} - z_2^{ij}), \tag{A8}$$

where the first term on the right hand side of eqn (A8) reflects trait divergence due to species sorting. The second term reflects trait divergence due to lineage sorting and the last term captures the trait differentiation within lineages. To estimate the magnitude of each of these components, we take the average of the absolute values of species sorting, lineage sorting and trait differentiation within lineages, respectively, obtained from eqn (A1) and eqn (A8), i.e.

$$
\begin{aligned}
&\frac{1}{2}\Big(\Big|\sum_{i=1}^{s} z_2^i(q_2^i - q_1^i)\Big| + \Big|\sum_{i=1}^{s} z_1^i(q_1^i - q_2^i)\Big|\Big),\\
&\frac{1}{2}\Big(\Big|\sum_{i=1}^{s} q_1^i \sum_{j=1}^{N_i} z_2^{ij}\Big(\frac{q_2^{ij}}{q_2^i} - \frac{q_1^{ij}}{q_1^i}\Big)\Big| + \Big|\sum_{i=1}^{s} q_2^i \sum_{j=1}^{N_i} z_1^{ij}\Big(\frac{q_1^{ij}}{q_1^i} - \frac{q_2^{ij}}{q_2^i}\Big)\Big|\Big),\\
&\frac{1}{2}\Big(\Big|\sum_{i=1}^{s} q_1^i \sum_{j=1}^{N_i} \frac{q_1^{ij}}{q_1^i}(z_2^{ij} - z_1^{ij})\Big| + \Big|\sum_{i=1}^{s} q_2^i \sum_{j=1}^{N_i} \frac{q_2^{ij}}{q_2^i}(z_1^{ij} - z_2^{ij})\Big|\Big).
\end{aligned}
\tag{A9}
$$

The first line in (A9) captures the average magnitude effect of species sorting, the second line the average magnitude effect of lineage sorting, and the last line the average effect of trait differentiation within lineages to the observed community trait divergence.

*Metric 2: The reaction norm approach*

The reaction norm equation, given by equation (A2) previously quantifies evolutionary and non-evolutionary contributions to community trait divergence using Community 1 as a reference. The reaction norm approach quantifying evolutionary and non-evolutionary components from Community 2 to Community 1 ($\Delta\bar{z}_{C_2\to C_1} = \bar{z}_1 - \bar{z}_2$) into the following components:

$$
\begin{aligned}
\Delta\bar{z}_{C_2\to C_1} = &\sum_{i=1}^{s} q_2^i(z_{21}^i - z_{22}^i) + \sum_{i=1}^{s} q_2^i(z_{12}^i - z_{22}^i)\\
&+ \sum_{i=1}^{s} q_2^i\big([z_{22}^i - z_{21}^i] - [z_{12}^i - z_{11}^i]\big)\\
&+ \sum_{i=1}^{s} z_{21}^i(q_1^i - q_2^i) + \sum_{i=1}^{s}(q_1^i - q_2^i)(z_{12}^i - z_{22}^i)\\
&+ \sum_{i=1}^{s}(q_1^i - q_2^i)\big([z_{22}^i - z_{21}^i] - [z_{12}^i - z_{11}^i]\big).
\end{aligned}
\tag{A10}
$$

The first term on the right hand side of eqn (A10) gives the trait divergence due to plasticity, the second term reflects genetic trait differentiation, the third term reflect genetic differentiation in plasticity, the fourth term captures trait divergence due to species sorting, and the last two terms reflect interaction components between species sorting and genetic trait differentiation and between species sorting and genetic differentiation in plasticity. We can calculate the average magnitude effects of each component by taking the average of the absolute values of the components given in eqn (A2) and (A10), i.e.

$$\begin{aligned}
&\frac{1}{2}\Big(\big|\sum_{i=1}^{s} q_1^i(z_{12}^i - z_{11}^i)\big| + \big|\sum_{i=1}^{s} q_2^i(z_{21}^i - z_{22}^i)\big|\Big)\\
&\frac{1}{2}\Big(\big|\sum_{i=1}^{s} q_1^i(z_{21}^i - z_{11}^i)\big| + \big|\sum_{i=1}^{s} q_2^i(z_{12}^i - z_{22}^i)\big|\Big)\\
&\frac{1}{2}\Big(\big|\sum_{i=1}^{s} q_1^i\big([z_{22}^i - z_{21}^i] - [z_{12}^i - z_{11}^i]\big)\big| + \big|\sum_{i=1}^{s} q_2^i\big([z_{22}^i - z_{21}^i] - [z_{12}^i - z_{11}^i]\big)\big|\Big)\\
&\frac{1}{2}\Big(\big|\sum_{i=1}^{s} z_{12}^i(q_2^i - q_1^i)\big| + \big|\sum_{i=1}^{s} z_{21}^i(q_1^i - q_2^i)\big|\Big)\\
&\frac{1}{2}\Big(\big|\sum_{i=1}^{s}(q_2^i - q_1^i)(z_{21}^i - z_{11}^i)\big| + \sum_{i=1}^{s}(q_1^i - q_2^i)(z_{12}^i - z_{22}^i)\big|\Big)\\
&\frac{1}{2}\Big(\big|\sum_{i=1}^{s}(q_2^i - q_1^i)\big([z_{22}^i - z_{21}^i] - [z_{12}^i - z_{11}^i]\big)\big| + \big|\sum_{i=1}^{s}(q_1^i - q_2^i)\big([z_{22}^i - z_{21}^i] - [z_{12}^i - z_{11}^i]\big)\big|\Big).
\end{aligned} \tag{A11}$$

In order of appearance the lines in (A11) represent magnitude effects for plasticity, genetic trait differentiation, genetic differentiation in plasticity, species sorting, species sorting × genetic trait differentiation and species sorting × genetic differentiation in plasticity.

*Metric 3: The Price-Reaction-Norm equation*

Equation (A3), previously given, can be used to quantify evolutionary and non-evolutionary contributions to community trait divergence going from Community 1 to Community 2. Similarly, evolutionary and non-evolutionary components to community trait divergence using Community 2 as a reference can be assessed as follows:

$$\begin{aligned}
\Delta\bar{z} = &\sum_{i=1}^{s} q_2^i \sum_{j=1}^{N_i} \frac{q_2^{ij}}{q_2^i}(z_{21}^{ij} - z_{22}^{ij}) + \sum_{i=1}^{s} q_2^i \sum_{j=1}^{N_i} z_{11}^{ij}\Big(\frac{q_1^{ij}}{q_1^i} - \frac{q_2^{ij}}{q_2^i}\Big)\\
&+ \sum_{i=1}^{s} q_2^i \sum_{j=1}^{N_i} \frac{q_2^{ij}}{q_2^i}(z_{12}^{ij} - z_{22}^{ij}) + \sum_{i=1}^{s} q_2^i \sum_{j=1}^{N_i} \frac{q_2^{ij}}{q_2^i}\big([z_{22}^{ij} - z_{21}^{ij}] - [z_{12}^{ij} - z_{11}^{ij}]\big)\\
&+ \sum_{i=1}^{s} z_{21}^i(q_1^i - q_2^i) + \sum_{i=1}^{s}(q_1^i - q_2^i)\sum_{j=1}^{N_i} z_{11}^{ij}\Big(\frac{q_1^{ij}}{q_1^i} - \frac{q_2^{ij}}{q_2^i}\Big)\\
&+ \sum_{i=1}^{s}(q_1^i - q_2^i)\sum_{j=1}^{N_i}\frac{q_2^{ij}}{q_2^i}(z_{12}^{ij} - z_{22}^{ij})\\
&+ \sum_{i=1}^{s}(q_1^i - q_2^i)\sum_{j=1}^{N_i}\frac{q_2^{ij}}{q_2^i}\big([z_{22}^{ij} - z_{21}^{ij}] - [z_{12}^{ij} - z_{11}^{ij}]\big).
\end{aligned} \tag{A12}$$

The first term on the right hand side of eqn (A12) gives the plasticity effect. The second term reflects lineage sorting, the third term gives the genetic trait differentiation within lineages, and the fourth term gives the trait differentiation in plasticity within lineages. The fifth term captures the species sorting, and the last three terms are interactions between species sorting and lineage sorting, species sorting and genetic trait differentiation within lineages and species sorting and genetic differentiation in plasticity within lineages. Taking the average of the absolute values given in eqn (A3) and (A12) result in the average magnitude effects of each component assessed by the Price-Reaction-Norm equation, i.e.

$$
\begin{aligned}
&\frac{1}{2}\Big(\Big|\sum_{i=1}^{s} q_1^i \sum_{j=1}^{N_i} \frac{q_1^{ij}}{q_1^i}(z_{12}^{ij} - z_{11}^{ij})\Big| + \Big|\sum_{i=1}^{s} q_2^i \sum_{j=1}^{N_i} \frac{q_2^{ij}}{q_2^i}(z_{21}^{ij} - z_{22}^{ij})\Big|\Big),\\
&\frac{1}{2}\Big(\Big|\sum_{i=1}^{s} q_1^i \sum_{j=1}^{N_i} z_{22}^{ij}\Big(\frac{q_2^{ij}}{q_2^i} - \frac{q_1^{ij}}{q_1^i}\Big)\Big| + \Big|\sum_{i=1}^{s} q_2^i \sum_{j=1}^{N_i} z_{11}^{ij}\Big(\frac{q_1^{ij}}{q_1^i} - \frac{q_2^{ij}}{q_2^i}\Big)\Big|\Big),\\
&\frac{1}{2}\Big(\Big|\sum_{i=1}^{s} q_1^i \sum_{j=1}^{N_i} \frac{q_1^{ij}}{q_1^i}(z_{21}^{ij} - z_{11}^{ij})\Big| + \Big|\sum_{i=1}^{s} q_2^i \sum_{j=1}^{N_i} \frac{q_2^{ij}}{q_2^i}(z_{12}^{ij} - z_{22}^{ij})\Big|\Big),\\
&\frac{1}{2}\Big(\Big|\sum_{i=1}^{s} q_1^i \sum_{j=1}^{N_i} \frac{q_1^{ij}}{q_1^i}([z_{22}^{ij} - z_{21}^{ij}] - [z_{12}^{ij} - z_{11}^{ij}])\Big| + \Big|\sum_{i=1}^{s} q_2^i \sum_{j=1}^{N_i} \frac{q_2^{ij}}{q_2^i}([z_{22}^{ij} - z_{21}^{ij}] - [z_{12}^{ij} - z_{11}^{ij}])\Big|\Big),\\
&\frac{1}{2}\Big(\Big|\sum_{i=1}^{s} z_{12}^i(q_2^i - q_1^i)\Big| + \Big|\sum_{i=1}^{s} z_{21}^i(q_1^i - q_2^i)\Big|\Big), \qquad\qquad \text{(A13)}\\
&\frac{1}{2}\Big(\Big|\sum_{i=1}^{s}(q_2^i - q_1^i) \sum_{j=1}^{N_i} z_{22}^{ij}\Big(\frac{q_2^{ij}}{q_2^i} - \frac{q_1^{ij}}{q_1^i}\Big)\Big| + \Big|\sum_{i=1}^{s}(q_1^i - q_2^i) \sum_{j=1}^{N_i} z_{11}^{ij}\Big(\frac{q_1^{ij}}{q_1^i} - \frac{q_2^{ij}}{q_2^i}\Big)\Big|\Big),\\
&\frac{1}{2}\Big(\Big|\sum_{i=1}^{s}(q_2^i - q_1^i) \sum_{j=1}^{N_i} \frac{q_1^{ij}}{q_1^i}(z_{21}^{ij} - z_{11}^{ij})\Big| + \Big|\sum_{i=1}^{s}(q_1^i - q_2^i) \sum_{j=1}^{N_i} \frac{q_2^{ij}}{q_2^i}(z_{12}^{ij} - z_{22}^{ij})\Big|\Big),\\
&\frac{1}{2}\Big(\Big|\sum_{i=1}^{s}(q_2^i - q_1^i) \sum_{j=1}^{N_i} \frac{q_1^{ij}}{q_1^i}([z_{22}^{ij} - z_{21}^{ij}] - [z_{12}^{ij} - z_{11}^{ij}])\Big|\\
&\qquad + \Big|\sum_{i=1}^{s}(q_1^i - q_2^i) \sum_{j=1}^{N_i} \frac{q_2^{ij}}{q_2^i}([z_{22}^{ij} - z_{21}^{ij}] - [z_{12}^{ij} - z_{11}^{ij}])\Big|\Big).
\end{aligned}
$$

order of appearance, (A13) gives the average magnitude effect of plasticity, lineage sorting, genetic trait differentiation, genetic differentiation in plasticity, species sorting, species sorting $\times$ lineage sorting, species sorting $\times$ genetic trait differentiation, species sorting $\times$ genetic differentiation in plasticity.

#### *Approach 3: Partitioning metrics for undirected trait change*

Next, we show how one can modify the community version of the Price equation, the reaction norm approach and the Price-Reaction-Norm equation to be independent of the reference chosen. This modification quantifies ecological and evolutionary components to pairwise observed community. Generally, we consider two spatially separated communities consisting of the same set of $s$ species. For the Price and Price-Reaction-Norm equation each species $i$ consists of $N_i$ genetic lineages. For the reaction norm and Price-Reaction-Norm equation, one assumes the construction of reaction norms for species, respectively for both species and genetic lineages. Here again, we denote the communities used in the pairwise comparison as Community 1 and Community 2.

##### *Metric 1: The Price equation*

The community version of the Price equation, given in equation (A1), quantifies species sorting, lineage sorting and trait change within lineages to community trait change going from Community 1 to Community 2. Previously, we described the contribution of species sorting, lineage sorting and trait change within lineage to community trait shift going from Community 2 to Community 1, given by equation (A8). The components in eqn (A1) and (A8) differ with respect to their multiplication with species trait values or relative abundances of either Community 1 ($z_1^i$, resp. $q_1^i$) or Community 2 ($z_2^i$, resp. $q_2^i$) depending on the reference. However, as stated in the main text for Approach 3 at the population level, one could instead use species average trait values and average relative abundances. This then modifies the community version of the Price equation to a spatial version independent of the reference choice, i.e.

$$\begin{aligned}\Delta\bar{z} = &\sum_{i=1}^{s}\Big(\frac{z_1^i + z_2^i}{2}\Big)(q_2^i - q_1^i) + \sum_{i=1}^{s}\Big(\frac{q_1^i + q_2^i}{2}\Big)\sum_{j=1}^{N_i}\Big(\frac{z_1^{ij} + z_2^{ij}}{2}\Big)\Big(\frac{q_2^{ij}}{q_2^i} - \frac{q_1^{ij}}{q_1^i}\Big)\\ &+ \sum_{i=1}^{s}\Big(\frac{q_1^i + q_2^i}{2}\Big)\sum_{j=1}^{N_i}\frac{1}{2}\Big(\frac{q_1^{ij}}{q_1^i} + \frac{q_2^{ij}}{q_2^i}\Big)(z_2^{ij} - z_1^{ij}),\end{aligned} \tag{A14}$$

where $z_k^i$ (resp. $z_k^{ij}$) is the average trait value of species $i$ (resp. lineage $j$ of species $i$) of Community $k$ and $q_k^i$ (resp. $q_k^{ij}$) is the relative abundance of species $i$ (resp. lineage $j$ of species $i$) in Community $k$. The first term on the right hand side of eqn (A14) reflects community trait divergence due to species sorting. The second term reflects trait divergence due to lineage sorting, and the last term reflects trait differentiation within lineages.

*Metric 2: The reaction norm approach*

As shown in the main text for Approach 3 the reaction norm approach cannot be straightforwardly modified to become independent of the reference point and still contain both main effects of plasticity and evolution and their interaction component. An alternative version of the reaction norm approach that is independent of the reference is given in a version of Ellner et al. (2011). This method assesses main effects of evolution and ecology and is independent of the reference. However, it does not include interaction components. We here modify the community version of the reaction norm approach given in eqn (A2) to become independent of the reference, by including the metric of Ellner et al. (2011). This is done by, in a first step, adding together the plasticity, constitutive evolution and evolution of plasticity component given in eqn (A2) and in eqn (A10) to give the observed trait divergence for each species $i$, i.e. $q_1^i(z_{22}^i - z_{11}^i)$ in the case of eqn (A2) and $q_2^i(z_{11}^i - z_{22}^i)$ in the case of eqn (A10). These terms are weighted by the species relative abundance of either Community 1 (eqn (A2)) or Community 2 (eqn (A10)). We instead replace this community-specific relative abundance by an average species relative abundance as given by Community 1 and Community 2, resulting in the following component: $\left(\frac{q_1^i+q_2^i}{2}\right)(z_{22}^i - z_{11}^i)$. Substituting eqn (15) from the main text into this component gives:

$$\begin{aligned}\left(\frac{q_1^i + q_2^i}{2}\right)(z_{22}^i - z_{11}^i) = &\sum_{i=1}^{s} \frac{1}{2}\left(\frac{q_1^i + q_2^i}{2}\right)(z_{12}^i - z_{11}^i + z_{22}^i - z_{21}^i) \\ &+ \sum_{i=1}^{s} \frac{1}{2}\left(\frac{q_1^i + q_2^i}{2}\right)(z_{21}^i - z_{11}^i + z_{22}^i - z_{12}^i).\end{aligned} \tag{A15}$$

In a second step, we add the interaction components in eqn (A2) or in eqn (A10) to the species sorting component, which results in the species sorting component of the Price equation (Govaert et al. 2016), i.e. $\sum_{i=1}^{s} z_{22}^{i}(q_2^i - q_1^i)$ for eqn (A2) and $\sum_{i=1}^{s} z_{11}^{i}(q_1^i - q_2^i)$ for eqn (A10). We then modify this component by using the average trait value of the species $(z_{11}^i + z_{22}^i)/2$ as opposed to the trait value in either Community 1 or Community 2. This results in a spatial community version of the reaction norm approach:

$$\Delta\bar{z} = \sum_{i=1}^{s} \frac{1}{2}\Big(\frac{q_1^i + q_2^i}{2}\Big)(z_{12}^i - z_{11}^i + z_{22}^i - z_{21}^i) + \sum_{i=1}^{s} \frac{1}{2}\Big(\frac{q_1^i + q_2^i}{2}\Big)(z_{21}^i - z_{11}^i + z_{22}^i - z_{12}^i) + \sum_{i=1}^{s} \Big(\frac{z_{11}^i + z_{22}^i}{2}\Big)(q_2^i - q_1^i), \tag{A16}$$

where $z_{kl}^i$ is the average trait value of species $i$ of Community $k$ at environmental condition $l$. The first term on the right hand side of eqn (A16) reflects community trait divergence due to the average effect of plasticity, the second term reflects average genetic trait differentiation and the last term reflects species sorting. It remains, however, unclear how eco-evolutionary interactions should be quantified.

*Metric 3: The Price-Reaction-Norm equation*

Constructing a spatial community version of the Price-Reaction-Norm equation combines both elements of the Price equation and of the reaction norm approach. Similarly as in the Price equation, we use average lineage and species trait values and average relative abundances. Similarly as in the reaction norm approach we incorporate eqn (15) of the main text by first adding the plasticity, genetic trait change and evolution of plasticity components together. Last, the interaction components (except for lineage sorting) are summed together with the species sorting component. This results in a spatial community version of the Price-Reaction-Norm equation that disentangles trait divergence between two spatially separated communities as follows:

$$\begin{aligned}
\Delta\bar{z} = &\sum_{i=1}^{s}\Big(\frac{q_1^i+q_2^i}{2}\Big)\sum_{j=1}^{N_i}\Big(\frac{z_{11}^{ij}+z_{22}^{ij}}{2}\Big)\Big(\frac{q_2^{ij}}{q_2^i}-\frac{q_1^{ij}}{q_1^i}\Big)\\
&+\sum_{i=1}^{s}\Big(\frac{q_1^i+q_2^i}{2}\Big)\sum_{j=1}^{N_i}\frac{1}{2}\Big(\frac{q_1^{ij}+q_2^{ij}}{2}\Big)(z_{12}^{ij}-z_{11}^{ij}+z_{22}^{ij}-z_{21}^{ij})\\
&+\sum_{i=1}^{s}\Big(\frac{q_1^i+q_2^i}{2}\Big)\sum_{j=1}^{N_i}\frac{1}{2}\Big(\frac{q_1^{ij}+q_2^{ij}}{2}\Big)(z_{21}^{ij}-z_{11}^{ij}+z_{22}^{ij}-z_{12}^{ij})\\
&+\sum_{i=1}^{s}(q_2^i-q_1^i)\sum_{j=1}^{N_i}\frac{1}{2}\Big(\frac{q_1^{ij}}{q_1^i}+\frac{q_2^{ij}}{q_2^i}\Big)\Big(\frac{z_{11}^{ij}+z_{22}^{ij}}{2}\Big)\\
&+\sum_{i=1}^{s}(q_2^i-q_1^i)\sum_{j=1}^{N_i}\Big(\frac{z_{11}^{ij}+z_{22}^{ij}}{2}\Big)\Big(\frac{q_2^{ij}}{q_2^i}-\frac{q_1^{ij}}{q_1^i}\Big).
\end{aligned}\tag{A17}$$

The first term on the right hand side of eqn (A17) gives the community trait divergence due to lineage sorting. The second term gives the average effect of plasticity, the third is the average genetic differentiation, the fourth is species sorting and the last is the interaction between species sorting and lineage sorting. A disadvantage of using eqn (15) to construct a spatial community version for the reaction norm approach and the Price-Reaction-Norm equation is that it remains unclear how the eco-evolutionary interactions should be quantified.

#### *Species gain and loss*

In the main text we assumed the same $s$ species were present in each of the $m$ spatially separated communities. However, this is an unlikely assumption in natural landscapes, or experimental treatments, and in many cases there will be only a subset of shared species between communities among a set of $m$ spatially separated communities. We can incorporate species gain and loss by subdividing the observed trait shift between a Community 1 and Community 2 across the shared and non-shared species:

$$\bar{z} = \sum_{i=1}^{s_2} q_2^i z_2^i - \sum_{i=1}^{s_1} q_1^i z_1^i = \sum_{i=1}^{s_c} q_2^i z_2^i - \sum_{i=1}^{s_c} q_1^i z_1^i + \sum_{i=s_c}^{s_2} q_2^i z_2^i - \sum_{i=s_c}^{s_1} q_1^i z_1^i, \tag{A18}$$

where $s_1$ and $s_2$ are the amount of species present in Community 1 and Community 2 and $s_c$ is the amount of shared species between Community 1 and Community 2. The first and second term on the right hand side of eqn (A18) is the observed trait shift between Community 1 and Community 2 across the shared species and the third and fourth term represent the species gain and loss. For the three suggested approaches, one can then use each of the three metrics (Price equation, reaction norm approach and Price-Reaction-Norm equation) to divide the trait shift among the shared species into evolutionary and non-evolutionary contributions as shown previously.

#### Appendix B: The Price-Reaction-Norm equation

Govaert et al. (2016) showed that the Price equation (as used in eqn (1) main text) could not separate phenotypic plasticity from genetic trait change and from evolution of plasticity within lineages, and confounds these processes in the within-lineage trait change component. The reaction norm approach performs better at this point (eqn (2) main text), but uses population means to calculate plasticity, constitutive evolution and evolution of plasticity, and therefore does not partition lineage sorting (Govaert et al. 2016). By combining these two approaches, a Price-Reaction-Norm equation can be constructed that is able to separate lineage sorting, genetic trait change and evolution of plasticity within lineages from phenotypic plasticity (Govaert et al. 2016). The Price-Reaction-Norm equation partitions the observed trait change between two time points in an asexually reproducing population consisting of $N$ genetic lineages, uniquely indexed by $j \in \{1, \ldots, N\}$ as follows:

$$\begin{aligned} \Delta\bar{z} = &\sum_{j=1}^{N} z_{22}^{j}(q_2^j - q_1^j) + \sum_{j=1}^{N} q_1^j (z_{21}^j - z_{11}^j) \\ &+ \sum_{j=1}^{N} q_1^j \left( [z_{22}^j - z_{21}^j] - [z_{12}^j - z_{11}^j] \right) + \sum_{j=1}^{N} q_1^j (z_{12}^j - z_{11}^j), \end{aligned} \tag{B1}$$

where $z_{kl}^j$ is the average trait value of genetic lineage $j$ at genetic state $k$ (i.e. sampled at time point $t_k$) and environmental state $l$ and $q_k^j$ is the relative abundance of genetic lineage $j$ at genetic state $k$ (i.e. sampled at time point $t_k$). The first term in the right hand side of eqn (B1) reflects lineage sorting, the second term captures genetic trait change within lineages, the third term captures evolution of plasticity, and the last term gives the plasticity component. This metric thus requires measurements of relative abundance for each genetic lineage at each time point (as in the Price equation) and of the lineage (average) trait values in the two environmental conditions (as in the reaction norm approach).

#### *Approach 1: Deviation from group mean*

Constructing a group mean in order to use the Price-Reaction-Norm (PRN) equation to partition trait divergence among multiple fragmented populations (which are composed of the same sets of genetic lineages, but see Appendix C when this assumption does not hold) into evolutionary and non-evolutionary components, requires calculating $z^j_{al}$ (i.e. the average trait value of lineage $j$ of the average population in environmental condition $l \in \{1, 2\}$), $z^j_{ka}$ (i.e. the average trait value of lineage $j$ of Population $k$ in the average environmental condition $a$), $z^j_{aa}$ (i.e. the average trait value of lineage $j$ of the average population in the average environmental condition), and $q^j_a$ (i.e. the average frequency of lineage $j$ across populations). The formulae to calculate these values are:

$$z^j_{al} = \frac{1}{m}\sum_{k=1}^{m} z^j_{kl}, \quad z^j_{ka} = \frac{z^j_{k1} + z^j_{k2}}{2}, \quad z^j_{aa} = \frac{1}{m}\sum_{k=1}^{m} z^j_{ka} \quad \text{and} \quad q^j_a = \frac{1}{m}\sum_{k=1}^{m} q^j_k. \tag{B2}$$

The PRN equation can then be used to partition the trait deviation from the group mean to each Population $k$ into the following components:

$$\begin{aligned}\Delta\bar{z}_{P_a \to P_k} = &\sum_{j=1}^{N} q^j_a (z^j_{al} - z^j_{aa}) + \sum_{j=1}^{N} z^j_{kl}(q^j_k - q^j_a) \\ &+ \sum_{j=1}^{N} q^j_a (z^j_{ka} - z^j_{aa}) + \sum_{j=1}^{N} q^j_a \left([z^j_{kl} - z^j_{ka}] - [z^j_{al} - z^j_{aa}]\right),\end{aligned} \tag{B3}$$

where the first term on the right hand side of eqn (B3) is the plasticity component, the second term is the trait deviation due to lineage sorting, the third component is the trait deviation due to genetic trait differentiation, and the last component is the trait deviation due to genetic differentiation in the plasticity response. Similarly as in the reaction norm approach, the plasticity component reflects the average plasticity response and not the absolute amount of plasticity of the population.

#### *Approach 2: Average of components in both directions*

The PRN equation separates differences in population trait values between two spatially separated populations into lineage sorting, genetic trait differentiation within lineages, genetic differentiation in plasticity within lineages and plasticity components (eqn (B1)). Eqn (B1) gives the partitioning in trait divergence using Population 1 as reference. Using Population 2 as reference, we can partition $\Delta\bar{z}_{2\rightarrow 1} = z_{11} - z_{22}$ into:

$$\begin{aligned}\Delta\bar{z}_{2\rightarrow 1} = &\sum_{j=1}^{N} z_{11}^{j}(q_1^j - q_2^j) + \sum_{j=1}^{N} q_2^j(z_{12}^j - z_{22}^j) \\ &+ \sum_{j=1}^{N} q_2^j\big([z_{22}^j - z_{21}^j] - [z_{12}^j - z_{11}^j]\big) + \sum_{j=1}^{N} q_2^j(z_{21}^j - z_{22}^j)\end{aligned} \tag{B4}$$

The terms on the right hand side of eqn (B4) reflect lineage sorting, genetic trait differentiation within lineages, genetic differentiation in plasticity within lineages, and plasticity, respectively. The absolute magnitude of each component is calculated by averaging the absolute value of each component of both directions, i.e.

$$\begin{aligned}&\frac{|\sum_{j=1}^{N} z_{22}^j(q_2^j - q_1^j)| + |\sum_{j=1}^{N} z_{11}^j(q_1^j - q_2^j)|}{2}, \\ &\frac{|\sum_{j=1}^{N} q_1^j(z_{21}^j - z_{11}^j)| + |\sum_{j=1}^{N} q_2^j(z_{12}^j - z_{22}^j)|}{2} \\ &\frac{|\sum_{j=1}^{N} q_1^j\big([z_{22}^j - z_{21}^j] - [z_{12}^j - z_{11}^j]\big)| + |\sum_{j=1}^{N} q_2^j\big([z_{22}^j - z_{21}^j] - [z_{12}^j - z_{11}^j]\big)|}{2}, \\ &\frac{|\sum_{j=1}^{N} q_2^j(z_{21}^j - z_{22}^j)| + |\sum_{j=1}^{N} q_2^j(z_{21}^j - z_{22}^j)|}{2}.\end{aligned} \tag{B5}$$

#### *Approach 3: Partitioning metrics for undirected trait change*

The Price-Reaction-Norm approach combines features of the Price equation and the reaction norm approach, and so our mathematical modification to a spatial version of this equation combines elements from the spatial extension of the Price equation and the reaction norm approach. Similarly as in the Price equation, depending whether we partition the trait change from Population 1 to Population 2 or vice versa, the lineage sorting component gives the difference between the relative abundances of the lineages of both populations multiplied with the trait value of the lineages of either Population 1 ($z^j_{11}$) or Population 2 ($z^j_{22}$). Multiplying the lineage sorting term instead with an average lineage trait value $(z^j_{11} + z^j_{22})/2$ as opposed to the lineage trait value of either Population 1 or 2, would result in a component of lineage sorting independent of the reference. The plasticity component and two evolutionary components in equation (B1) and (B4) are weighted by the relative abundance of lineage j of Population 1 or Population 2 depending on whether Population 1 or Population 2 is taken as a reference. Multiplying instead with an average relative abundance of lineage $j$ (i.e. $(q^j_1 + q^j_2)/2$) gives the following modified version of the Price-Reaction-Norm equation:

$$\begin{aligned}\Delta\bar{z} = &\sum_{j=1}^{N}\Big(\frac{z^j_{11}+z^j_{22}}{2}\Big)(q^j_2 - q^j_1) + \sum_{j=1}^{N}\Big(\frac{q^j_1+q^j_2}{2}\Big)(z^j_{21} - z^j_{11}) \\ &+ \sum_{j=1}^{N}\Big(\frac{q^j_1+q^j_2}{2}\Big)([z^j_{22} - z^j_{21}] - [z^j_{12} - z^j_{11}]) + \sum_{j=1}^{N}\Big(\frac{q^j_1+q^j_2}{2}\Big)(z^j_{12} - z^j_{11}).\end{aligned} \tag{B6}$$

The last three components on the right hand side of eqn (B6) still depend on the choice of reference. Similarly as in the reaction norm approach, we did not find a way to spatially modify the plasticity, genetic trait differentiation and genetic differentiation in plasticity to become independent of the reference point. However, the sum of the plasticity, genetic trait differentiation and genetic differentiation in plasticity components adds up to an average-abundance weighted trait change of lineage $j$; i.e. $\sum_{j=1}^{N}(\frac{q^j_1+q^j_2}{2})(z^j_{22} - z^j_{11})$. We previously suggested the use of eqn (15) (eqn (12) in Ellner et al. 2011) as a spatial version of the reaction norm approach. We thus here, modify eqn (A17) by substituting $\Delta z^j = z^j_{22} - z^j_{11}$ by eqn (15). This allows to partition $z^j_{22} - z^j_{11}$ into an average plasticity and average genetic trait differentiation effect. This results in a spatial version of the Price-Reaction-Norm equation, given by the following equation:

$$\begin{aligned}\Delta\bar{z} = &\sum_{j=1}^{N}\Big(\frac{z^j_{11}+z^j_{22}}{2}\Big)(q^j_2 - q^j_1) + \sum_{j=1}^{N}\Big(\frac{q^j_1+q^j_2}{2}\Big)\Big[\frac{(z^j_{12} - z^j_{11}) + (z^j_{22} - z^j_{21})}{2}\Big] \\ &+ \sum_{j=1}^{N}\Big(\frac{q^j_1+q^j_2}{2}\Big)\Big[\frac{(z^j_{21} - z^j_{11}) + (z^j_{22} - z^j_{12})}{2}\Big],\end{aligned} \tag{B7}$$

where the first term on the right hand side of eqn (B7) represents the trait divergence due to lineage sorting, the second term reflects the average plasticity effect, and the last term gives the average effect of genetic trait differentiation. While this is a suitable decomposition of trait change in space in a way that is independent of the choice of reference, it has the drawback that genetic trait differentiation of plasticity cannot be quantified.

#### Appendix C: Genetic lineage gain and loss

The assumption made in the main text for the Price equation and the Price-Reaction-Norm equation is that the spatially separated populations consist of the same $N$ genetic lineages, uniquely indexed as $j \in \{1, \ldots, N\}$. This assumption may not hold for natural populations. However, there might be some overlap of lineages between subsets of populations (as illustrated in Fig. C1). In order to apply the Price equation and Price-Reaction-Norm equation to this type of data (i.e. some overlap of genetic lineages among the spatially separated populations, but not all lineages are the same) you can separate the observed trait change among the shared and non-shared sets of genetic lineages. For example, for two populations (Population $k$ and Population $r$) that share $N_c$ genetic lineages and in which the shared lineages are indexed first, this would become:

$$\bar{z} = \sum_{j=1}^{N_r} q_r^j z_r^j - \sum_{j=1}^{N_k} q_k^j z_k^j = \sum_{j=1}^{N_c} q_r^j z_r^j - \sum_{j=1}^{N_c} q_k^j z_k^j + \sum_{j=N_c}^{N_r} q_r^j z_r^j - \sum_{j=N_c}^{N_k} q_k^j z_k^j. \tag{C1}$$

In eqn (C1) $z_k^j$ and $q_k^j$ refer to the trait value and relative abundance of genetic lineage $j$ of Population $k$. The first two terms on the right hand side of eqn (C1) represent the trait change from Population $k$ to Population $r$ across the shared genetic lineages. The last two term represent the trait change that is due to lineages present in Population $r$ but not in Population $k$ and vice versa. Note that when none of the lineages are shared the first two terms on the right hand side of eqn (C1) would equal zero. We next detail how the three approaches proposed in the main text apply to $m$ spatially separated populations where some but not all genetic lineages are shared. We only show the modifications for the Price and Price-Reaction-Norm equation, as the reaction norm approach does not require information on genetic lineages.

#### *Approach 1: Deviation from group mean*

We thus consider a set of $m$ spatially separated populations, in which genetic lineages are shared among some sets of populations. The group mean can then be constructed as follows:

$$z_a^j = \frac{\sum_{k=1}^m c_{kj} z_k^j}{\sum_{k=1}^m c_{kj}} \quad \text{and} \quad q_a^j = \frac{1}{m}\sum_{k=1}^m q_k^j, \tag{C2}$$

where $z_a^j$ is the average trait value of genetic lineage $j$, $c_{kj} = 1$ when genetic lineage $j$ is present in Population $k$ and $c_{kj} = 0$ otherwise, and $q_a^j$ represents the average relative abundance of genetic lineage $j$. By default, the group mean will contain all genetic lineages among the set of $m$ spatially separated populations (Fig. C1), and there will thus always be at least one genetic lineage shared between the group mean and one of the $m$ populations. Because the Price equation and Price-Reaction-Norm equation can only be applied on the trait change across the shared set of lineages, we first separate the observed trait deviation from the group mean to each Population $k$ across the shared and non-shared genetic lineages as in eqn (C1). However, because no genetic lineage can be gained from the group mean to a Population $k$, we do not have a genetic lineage gain component, i.e.

$$\Delta \bar{z}_{P_a \to P_k} = \sum_{j=1}^{N_c^k} q_k^j z_k^j - \sum_{j=1}^{N_c^k} q_a^j z_a^j - \sum_{j=N_c^k}^{N_a} q_a^j z_a^j. \tag{C3}$$

In eqn (C3) the first two terms on the right hand side are summed over the shared genetic lineages where $N_c^k$ denotes the total amount of shared genetic lineages of Population $k$ with the group mean. The last term contains those genetic lineages that are only present in the group mean and not in Population $k$. This term reflects a loss of genetic lineages from a regional lineage pool (i.e. the group mean) to a local site. We can then for example, use the Price equation (eqn (1) main text) to quantify lineage sorting and trait deviation within lineages to the trait deviation from the group mean to Population $k$ across the shared set of genetic lineages, i.e.

$$\Delta \bar{z}_{P_a \to P_k} = \sum_{j=1}^{N_c^k} z_k^j (q_k^j - q_a^j) + \sum_{j=1}^{N_c^k} q_a^j (z_k^j - z_a^j) - \sum_{j=N_c}^{N_a} q_a^j z_a^j. \tag{C4}$$

Similarly, we could have used the Price-Reaction-Norm approach (eqn (B1)) in Appendix B) on the set of shared genetic lineages. This would partition the trait deviation from the group mean to a Population $k$ as follows:

$$\begin{aligned}\Delta\bar{z}_{P_a\rightarrow P_k} &= \sum_{j=1}^{N_c^k} q_a^j(z_{al}^j - z_{aa}^j) + \sum_{j=1}^{N_c^k} z_{kl}^j(q_k^j - q_a^j)\\ &+ \sum_{j=1}^{N_c^k} q_a^j(z_{ka}^j - z_{aa}^j) + \sum_{j=1}^{N_c^k} q_a^j\left([z_{kl}^j - z_{ka}^j] - [z_{al}^j - z_{aa}^j]\right) - \sum_{j=N_c}^{N_a} q_a^j z_{aa}^j.\end{aligned} \tag{C5}$$

In eqn (C5), $z_{al}^j$ equals the average trait value of genetic lineage $j$ in environmental condition $l$. This average is taken across the sites genetic lineage $j$ is present. $z_{ka}^j$ is the trait value of genetic lineage $j$ of Population $k$ at the average environmental condition $a$. $z_{aa}^j$ equals the average trait value of genetic lineage $j$ at the average environmental condition $a$. Last, $q_a^j$ equals the average abundance of genetic lineage $j$. Specifically, these are calculated as follows:

$$z_{al}^j = \frac{\sum_{k=1}^m c_{kj} z_{kl}^j}{\sum_{k=1}^m c_{kj}}, \quad z_{ka}^j = \frac{z_{k1}^j + z_{k2}^j}{2}, \quad z_{aa}^j = \frac{\sum_{k=1}^m c_{kj} z_{ka}^j}{\sum_{k=1}^m c_{kj}} \quad \text{and} \quad q_a^j = \frac{1}{m}\sum_{k=1}^m q_k^j, \tag{C6}$$

where $c_{kj} = 1$ when genetic lineage $j$ is present in Population $k$, and $c_{kj} = 0$ otherwise. $z_{ka}^j$ can only be calculated for those populations in which genetic lineage $j$ occurs. In eqn (C5), the first term on the left hand side reflects the average plasticity response to environmental condition $l$. The second term captures the trait deviation from the group mean to lineage sorting across the shared lineages. The third and fourth term capture the genetic trait differentiation and the genetic differentiation in plasticity across the shared lineages to the trait deviation from the group mean to Population $k$. The last term reflects that part of the trait deviation that is due to genetic lineage loss from the group mean to Population $k$.

#### *Approach 2: Average of components in both directions of comparison*

In the next approach, evolutionary and non-evolutionary contributions are calculated between pairs of populations (e.g. Population $k$ and $r$). Thus, among a set of $m$ spatially separated populations, one calculates evolutionary and non-evolutionary contributions to trait shift from Population $k$ to Population $r$ and vice versa. Of these contributions, the average of the absolute values are taken and magnitudes of evolutionary and non-evolutionary contributions are obtained. Because the Price equation and the Price-Reaction-Norm approach can only be applied on the set of shared genetic lineages, we first need to separate the trait shift between pairs of populations into a set of shared and non-shared genetic lineages. Assuming a trait shift from Population $k$ to Population $r$ this gives:

$$\Delta \bar{z}_{P_k \to P_r} = \sum_{j=1}^{N_c} q_r^j z_r^j - \sum_{j=1}^{N_c} q_k^j z_k^j + \sum_{j=N_c}^{N_r} q_r^j z_r^j - \sum_{j=N_c}^{N_k} q_k^j z_k^j. \tag{C7}$$

Similarly, for a trait shift from Population $r$ to Population $k$ this gives:

$$\Delta \bar{z}_{P_r \to P_k} = \sum_{j=1}^{N_c} q_k^j z_k^j - \sum_{j=1}^{N_c} q_r^j z_r^j + \sum_{j=N_c}^{N_k} q_k^j z_k^j - \sum_{j=N_c}^{N_r} q_r^j z_r^j. \tag{C8}$$

Here $N_c$ is the set of shared lineages between Population $k$ and $r$, starting from index $j = 1$. In both eqn (C7) and (C8) the first two terms on the right hand side give the trait shift between Population $k$ and $r$ across the shared genetic lineages between the two populations. The last two terms in eqn (C7) and (C8) represent that part of the trait shift that is due to non-sharing genetic lineages. Logically, the genetic lineages lost from Population $k$ to $r$ (last term eqn (C7)) are gained going from Population $r$ to $k$ (second last term in eqn (C8)), and vice versa. Hence, $\sum_{j=N_c}^{N_r} q_r^j z_r^j - \sum_{j=N_c}^{N_k} q_k^j z_k^j = -(\sum_{j=N_c}^{N_r} q_r^j z_r^j - \sum_{j=N_c}^{N_k} q_k^j z_k^j)$ and thus the sum of both terms from eqn (C7) and (C8) equal each other in magnitude but are opposite in sign and capture that both populations have the same degree of trait shift due to non-shared lineages. Using the Price equation and the proposed modification from Approach 2, we can calculate the average of the absolute values of the evolutionary and non-evolutionary components across the set of shared lineages, i.e.

$$\begin{aligned} &\frac{|\sum_{j=1}^{N_c} z_r^j (q_r^j - q_k^j)| + |\sum_{j=1}^{N_c} z_k^j (q_k^j - q_r^j)|}{2}, \\ &\frac{|\sum_{j=1}^{N_c} q_k^j (z_r^j - z_k^j)| + |\sum_{j=1}^{N_c} q_r^j (z_k^j - z_r^j)|}{2}, \\ &|\sum_{j=N_c}^{N_k} q_k^j z_k^j - \sum_{j=N_c}^{N_r} q_r^j z_r^j|. \end{aligned} \tag{C9}$$

The first line in (C9) gives the magnitude of lineage sorting between Population $k$ and $r$. The second line gives the magnitude of trait differentiation between Population $k$ and $r$, and the last line gives the magnitude of trait differentiation due to non-shared lineages between Population $k$ and $r$. Note if there are no overlapping lineages between the two populations, then we only have the last line in (C9).

Similarly, we could substitute the first two terms of eqn (C7) and (C8) with the Price-Reaction-Norm equation, and calculate the average of the absolute values of the evolutionary and non-evolutionary components across the set of shared lineages, i.e.

$$\begin{aligned}
&\frac{|\sum_{j=1}^{N_c} z_{r2}^j (q_r^j - q_k^j)| + |\sum_{j=1}^{N_c} z_{k1}^j (q_k^j - q_r^j)|}{2}, \\
&\frac{|\sum_{j=1}^{N_c} q_k^j (z_{r1}^j - z_{k1}^j)| + |\sum_{j=1}^{N_c} q_r^j (z_{k2}^j - z_{r2}^j)|}{2} \\
&\frac{|\sum_{j=1}^{N_c} q_k^j ([z_{r2}^j - z_{r1}^j] - [z_{k2}^j - z_{k1}^j])| + |\sum_{j=1}^{N_c} q_r^j ([z_{r2}^j - z_{r1}^j] - [z_{k2}^j - z_{k1}^j])|}{2}, \\
&\frac{|\sum_{j=1}^{N_c} q_k^j (z_{k2}^j - z_{k1}^j)| + |\sum_{j=1}^{N_c} q_r^j (z_{r1}^j - z_{r2}^j)|}{2}, \\
&|\sum_{j=N_c}^{N_k} q_k^j z_{k1}^j - \sum_{j=N_c}^{N_r} q_r^j z_{r2}^j|.
\end{aligned} \tag{C10}$$

In eqn (C10), Population $k$ is assumed to originate from a site with environmental condition 1, and Population $r$ is assumed to orginate from a site with environmental condition 2. The first line of eqn (C10) captures the magnitude of lineage sorting, the second term captures the magnitude of genetic trait differentiation, the third term captures the genetic differentiation in plasticity, the fourth term reflect the average magnitude of plasticity, and the last reflects the average magnitude of trait differentiation due to non-shared lineages between the two populations.

#### *Approach 3: Partitioning metrics for undirected trait change*

In the last approach we mathematically modified partitioning metrics to account for undirected trait change between pairs of populations. Similarly as in Approach 2, we can write the trait shift in sums of shared versus non-shared genetic lineages between pairs of populations (say

Population $k$ and $r$) as in eqn (C1). We can then use the spatial version of the Price equation given in eqn (14) from the main text on the first two terms of eqn (C1) to give the contribution of lineage sorting and trait differentiation between the two populations, i.e.

$$\Delta\bar{z} = \sum_{j=1}^{N_c}\left(\frac{z_k^j + z_r^j}{2}\right)(q_r^j - q_k^j) + \sum_{j=1}^{N_c}\left(\frac{q_k^j + q_r^j}{2}\right)(z_r^j - z_k^j) + \sum_{j=N_c}^{N_r} q_r^j z_r^j - \sum_{j=N_c}^{N_k} q_k^j z_k^j. \tag{C11}$$

Depending whether we chose Population $k$ or Population $r$ as a reference, the last two terms in eqn (C11) will change in sign but not in magnitude. These terms reflect the trait shift between the two populations due to different lineages being present.

For the Price-Reaction-Norm equation, we can use eqn (B7) in Appendix B to partition the trait differentiation between Population $k$ and $r$ as follows:

$$\begin{aligned}\Delta\bar{z} &= \sum_{j=1}^{N_c}\left(\frac{z_{k1}^j + z_{r2}^j}{2}\right)(q_r^j - q_k^j) + \sum_{j=1}^{N_c}\left(\frac{q_k^j + q_r^j}{2}\right)\left[\frac{(z_{k2}^j - z_{k1}^j) + (z_{r2}^j - z_{r1}^j)}{2}\right] \\ &+ \sum_{j=1}^{N_c}\left(\frac{q_k^j + q_r^j}{2}\right)\left[\frac{(z_{r1}^j - z_{k1}^j) + (z_{r2}^j - z_{k2}^j)}{2}\right] + \sum_{j=N_c}^{N_r} q_r^j z_r^j - \sum_{j=N_c}^{N_k} q_k^j z_k^j.\end{aligned} \tag{C12}$$

The first term on the right hand side of eqn (C12) captures the trait divergence due to lineage sorting, the second term reflects the average plasticity term, and the last term gives the average effect of genetic trait differentiation, all across the shared genetic lineages. The last term then reflects the trait shift between the two populations due to different lineages present.

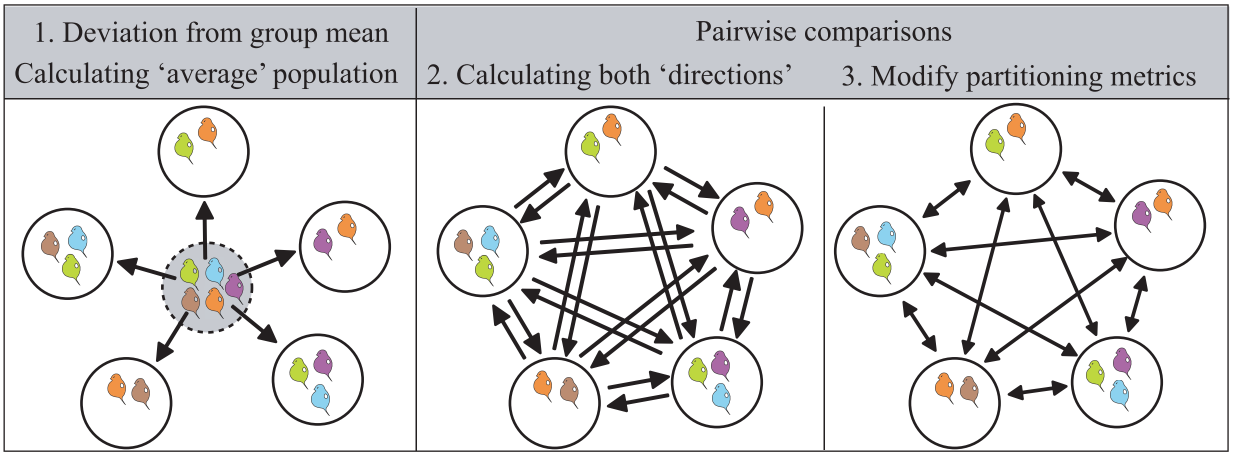


Figure C1: Visualization of a situation in which different genetic lineage of 5 *Daphnia* clones (depicted in brown, blue, orange, purple and green) are present across 5 spatial locations.

#### Appendix D: Constructing group mean reaction norm approach

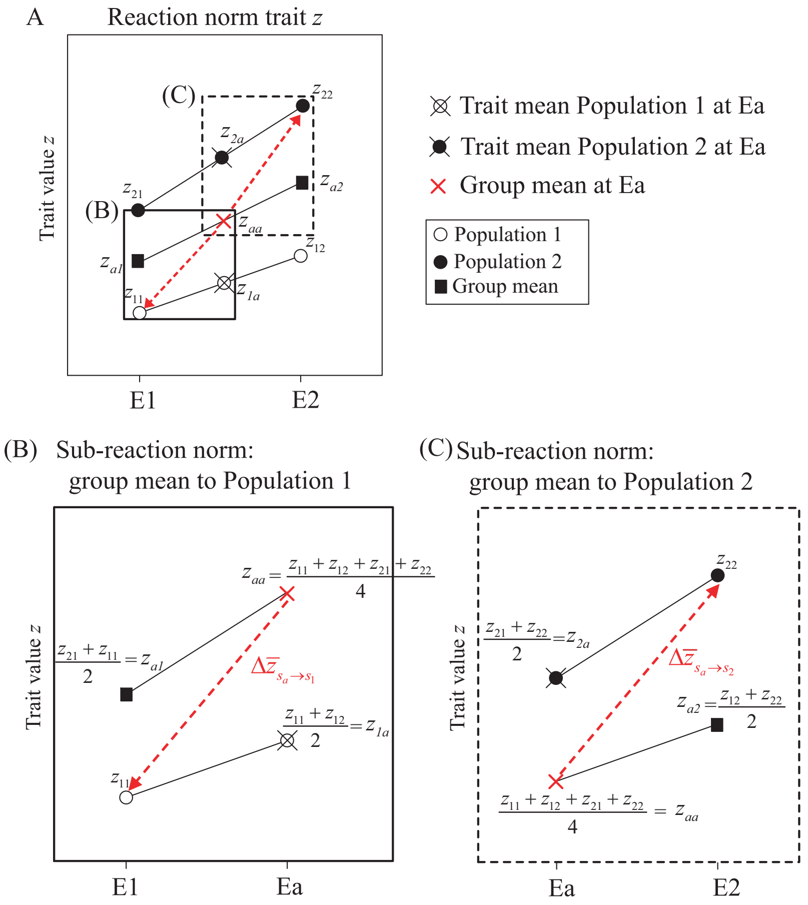


Figure D1: (A) Visualisation of the construction of the group mean (filled squares) between two spatially separated populations (Population 1: unfilled circles; Population 2: filled circles) when using the reaction norm approach. $z_{kl}$ equals the trait mean of Population $k$ in environmental condition $l$. A group mean is constructed by calculating $z_{ka}$ (the average trait value of Population $k$ in the average environmental condition $a$), $z_{al}$ (the average trait value of the average population in environmental condition $l$) and $z_{aa}$ (the average trait value of the average population in the average environmental condition) as given in the main text (detailed in eqn (5)), but here applied to a set of 2 spatially separated populations (i.e. $k \in \{1, 2\}$). Panel (B) and (C) zoom in on subparts of the reaction norm, where one separates the shift in trait mean from the group mean in the 'average' environmental condition (Ea) to (B) Population 1 at environmental condition E1 and (C) Population 2 at environmental condition E2.

### Appendix E: Contributions of plasticity and evolution to expected phenotypic responses of *Chamaecrista fasciculata*, a North American Prairie plant (Etterson and Shaw (2001))

Etterson and Shaw (2001) determined the phenotypic response of the North American Prairie plant *Chamaecrista fasciculata* to global warming. They performed a transplant experiment with three populations sampled from a gradient from cold and wet environments to warm and arid environments: Minnesota (MN), Kansas (KS), and Oklahoma (OK). Because it was assumed that the MN population would experience similar climatic conditions as KS and OK as climate change proceeds (Etterson and Shaw 2001), we can infer a direction of trait change in this study. We decided to treat the MN population as ancestral population and the KS and OK populations as descendent populations. This implies a directed trait change from the MN population to the KS and OK populations. This study thus gives an example of a spatial study system in which a directed trait change can be assumed.

From the measurements taken during the transplant experiment, reaction norms can be constructed, which allows the use of the reaction norm approach. We use the reaction norm approach on two traits - the number of leaves and leaf thickness ($m^2/g$; leaf area divided by dry leaf weight) - with values obtained from the reciprocal transplant experiment, to determine the contributions of plasticity, constitutive evolution and evolution of plasticity to the expected phenotypic response of this population to climate change (Fig. E1). Phenotypic plasticity was found to be more important than evolutionary change for the changes in leaf number, while evolutionary trait change was more important than plasticity for the changes in leaf thickness. The evolutionary response was dominated by evolution of plasticity, whereas evolution of mean trait value contributed little to change in leaf thickness, and this for both trait change from MN to KS and from MN to OK (Fig. E1C,F). From the reaction norm approach, we can thus derive which processes are important contributors to the expected trait change corresponding to a shift from cold and wet environments to warm and arid environments.

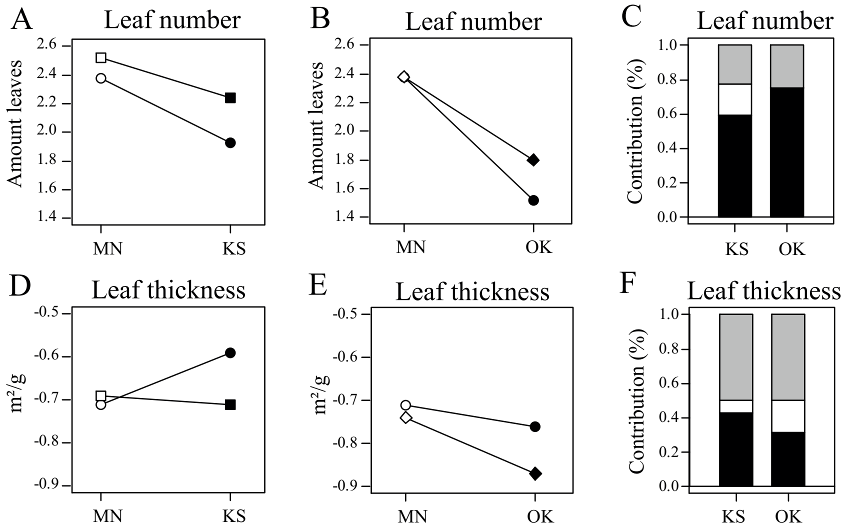


Figure E1: Graphical representation of the reaction norms for (A-B) leaf number and (D-E) leaf thickness of the three native populations of *Chamaecrista fasciculata* from Minnesota (MN), Kansas (KS) and Oklahoma (OK) from which individuals were transplanted into field sites in MN, KS and OK. (C,F) Relative contributions of plasticity (black), constitutive evolution (white) and evolution of plasticity (grey) are calculated using the reaction norm approach for trait change from MN to KS and from MN to OK. Data values were extracted from Figure 3 in Etterson & Shaw (2001).

### Appendix F: Application to empirical examples

In the main text, we wanted to assess whether *A. maculatum* populations differed in their ecological and evolutionary contributions depending on whether they originated from a pond where the predator *A. opacum* was present or absent (data from Urban 2008). We used the reaction norm approach described in Approach 1 and quantified ecological and evolutionary contributions to trait deviations from a group mean to each of the *A. maculatum* populations. These contributions can assess the repeatability in the contributions of plasticity and evolution to trait shift from an average or local predator-free pond to each *A. maculatum* population inhabiting predator ponds, and vice versa (Example 1A) or between pairs of populations where each pair consists of a population from predator-free and predator-present pond (Example 1B).

#### *Example 1A: Repeatability in the contributions of plasticity and evolution to trait shift from an average or local predator-free pond to each A. maculatum population inhabiting predator ponds, and vice versa*

To test for the repeatability in the contributions of plasticity and evolution, we ask the following set of questions:

(1a) Do we detect similar contributions of plasticity and evolution if all *A. maculatum* populations inhabiting predator ponds descended from an average predator-free pond?

(1b) Do we detect similar contributions of plasticity and evolution if all *A. maculatum* populations inhabiting predator-free ponds descended from an average predator pond?

(2a) Do we detect similar contributions of plasticity and evolution if all *A. maculatum* populations inhabiting predator ponds descended from one of the existing predator-free ponds?

(2b) Do we detect similar contributions of plasticity and evolution if all *A. maculatum* populations inhabiting predator-free ponds descended from one of the existing predator-present ponds?

For the first two questions we need to first calculate an average *A. maculatum* population inhabiting a predator-free (Question 1a) and predator-present (Question 1b) pond, and use this population as a reference in subsequent calculations of eco-evolutionary contributions to observed trait shifts in prey body mass. Partitioning the trait shift from the average predator-free population ($A$) to each of the predator-present populations using the reaction norm approach translates into the following:

$$\Delta z_{A \to k} = (z_{A1} - z_{A0}) + (z_{k0} - z_{A0}) + ([z_{k1} - z_{k0}] - [z_{A1} - z_{A0}]), \tag{F1}$$

in which $z_{A1}$ (resp. $z_{A0}$) represent the trait value of the average predator-free *A. maculatum* population in the predator-kairomone (resp. control) condition, and $z_{k1}$ (resp. $z_{k0}$) is the trait value of a predator-present population $k$ in the predator-kairomone (resp. control) condition. The first term on the right hand side of eqn (F1) is the plasticity component of the average predator-free population, the second term is the constitutive evolution component and the third term is the evolution of plasticity component. Note that as the first term is the plasticity component of the average predator-free population, it will have the same value across the different predator populations. Hence, these populations will thus only differ in their constitutive and evolution of plasticity component. Similarly, we can calculate the contributions of plasticity, constitutive evolution and evolution of plasticity to the trait shift from the average predator-present population to each of the predator-free *A. maculatum* populations using the following equation:

$$\Delta z_{P \to k} = (z_{P0} - z_{P1}) + (z_{k1} - z_{P1}) + ([z_{k0} - z_{k1}] - [z_{P0} - z_{P1}]). \tag{F2}$$

In this equation, $z_{P1}$ (resp. $z_{P0}$) represent the trait value of the average predator *A. maculatum* population in the predator-kairomone (resp. control) condition. Again, as the first term on the right hand side of eqn (F2) reflects the plasticity component of the average predator-present population, it will have the same value across the predator-free populations.

For questions (2a) and (2b), we can choose one of the *A. maculatum* populations from either an existing predator-free (Question 2a) or predator-present (Question 2b) pond, and use this population as a reference in subsequent calculations of eco-evolutionary contributions. For both sets of questions we then use the reaction norm approach to calculate contributions of plasticity, constitutive evolution and evolution of plasticity to one of the existing predator-free populations (noted with $r$) to each of the predator-present populations (noted with $k$):

$$\Delta z_{r\to k} = (z_{r1} - z_{r0}) + (z_{k0} - z_{r0}) + ([z_{k1} - z_{k0}] - [z_{r1} - z_{r0}]), \tag{F3}$$

or to one of the existing predator-present populations to each of the predator-free populations:

$$\Delta z_{k\to r} = (z_{k0} - z_{k1}) + (z_{r1} - z_{k1}) + ([z_{r0} - z_{r1}] - [z_{k0} - z_{k1}]). \tag{F4}$$

In both equations (F3) and (F4), $z_{r1}$ (resp. $z_{r0}$) is the trait value of the $r$th predator-free *A. maculatum* population in the predator-kairomone (resp. control) condition, and $z_{k1}$ (resp. $z_{k0}$) is the trait value of the $k$th predator-present *A. maculatum* population in the predator-kairomone (resp. control) condition.

Absolute contributions of constitutive evolution were in the same direction for trait shifts from the average predator-present population to all predator-free populations (Fig. F1). This was, however, not the case for the absolute contributions of constitutive evolution to trait shift from the average predator-free population to all of the predator-present populations, or in general for the evolution of plasticity contributions. This shows that the repeatability in the sign of eco-evolutionary directions may depend on the strength of the selection pressure (e.g. strong response of constitutive evolution reduces mean prey size when selection shifts from no predation to predation). When calculating relative contributions (Fig. F2), we found that plasticity was the stronger contributor to the expected trait shifts. Both absolute and relative contributions of constitutive evolution and evolution of plasticity varied among both the *A. maculatum* populations originating from predator-free ponds and descending from an average predator-present population as for the *A. maculatum* populations originating from predator-present ponds and descending from an average predator-free population.

For the second set of questions (Question 2a and 2b), we treated each of the existing ponds as the ancestral population. Comparing the absolute contributions of constitutive evolution and evolution of plasticity among the different reference populations used (represented by the different colors in Figure F3), shows that the descendent populations differ from one another. But using a different reference population only results in a horizontal and/or vertical shift of the absolute contributions of these two components (Fig. F3). This also illustrates that choosing a different reference population will change contributions of the evolutionary and non-evolutionary components, but the results will still reflect qualitative differences between the descendant populations. Relative contributions can be visualized either using heatmaps (Fig. F4) or via triangle plots (Fig. F5). Overall, we found that plasticity had the largest relative contributions to change in prey body mass both for going from all predator-free *A. maculatum* populations to predator-present *A. maculatum* populations and from all predator-present *A. maculatum* populations to predator-free *A. maculatum* populations (Fig. F4A,B). Relative contributions of constitutive evolution vary more for the predator-free to predator-present direction (Fig. F4C,D), while relative contributions of evolution of plasticity vary more for the predator-present to predator-absent direction (Fig. F4E,F and Fig. F5).

##### *Example 1B: Contributions of ecological and evolutionary processes to trait shifts between pairs of A. maculatum populations*

The data set collected by Urban (2008) can also be used to evaluate the overall contribution of plasticity, genetic trait differentiation and genetic differentiation in plasticity between a *A. maculatum* population originating from a pond in which the predator *A. opacum* is present and a *A. maculatum* population originating from a pond in which the predator *A. opacum* is absent. While this question is very similar to the previously discussed Question 2a and 2b, it crucially differs from it in the assumption of the reference. In Question 2a and 2b, we assumed there was a trait shift from one of the predator-free or predator-present to one of the predator-present or predator-free populations, while in this example, such an assumption is not made. We, instead, quantify contributions of plasticity, genetic trait differentiation and genetic differentiation in plasticity among pairwise combinations of predator-present and predator-free *A. maculatum* populations using the reaction norm approach as described in Approach 2. In this approach, absolute values of eco-evolutionary contributions to trait shift from predator-present to predator-free and vice versa are averaged, within each pair. Some populations had consistently smaller contributions of one of the three processes, indicating that these processes may depend on site identity (Fig. F6-F7). For example, the predator-present population *canis* overall showed similar contributions among all pairwise comparisons with predator-free *A. maculatum* populations, except for the predator-free *p7* and *p24* population (F6). For these populations, lower contributions of plasticity and genetic differentiation in plasticity and higher contributions of genetic trait differentiation were found. This means that in these populations other local selection pressures potentially acted on the trait values of those particular populations resulting in different prey body mass. Moreover, contributions in pairs of which the population *canis* was present were clearly distinct from contributions of other pairs were this populations was not present (Fig. F7). From Figure F7, we can detect subsets of combinations that occupy similar or distinct subspaces of eco-evolutionary contributions. The mechanisms to explain and predict whether populations will have higher or lower relative contributions of a specific process or why certain populations show similar eco-evolutionary contributions are interesting future research questions.

#### *Example 3: Are spatial and temporal trait divergence structured by the same processes? – A case study in Daphnia*

In example 3 in the main text, we found a similar range in the absolute contributions of plasticity, genetic trait differentiation and genetic differentiation in plasticity between (sub)populations of *Daphnia magna* that experienced similar fish predation pressure comparing a temporal (Cousyn et al. 2001) and a spatial (De Meester 1996) setting. We determined this using the reaction norm approach on trait deviations from the group mean. We here show that this result is observed independent of the approach used. Thus, using the reaction norm approach as given in Approach

2 (eqn (11), (12) and (13) main text) and Approach 3 (eqn (15), we can quantify the absolute and relative contributions of plasticity, genetic trait differentiation and genetic differentiation in plasticity between pairs of *Daphnia* (sub)populations (Fig. F8). We also used bootstrapping to construct confidence intervals around the estimated values, by sampling the data 1000 times and recalculating contributions of plasticity, genetic trait differentiation and genetic differentiation in plasticity. Similarly as in the main text, we did not find differences between the absolute contributions of pairwise comparisons between the fish-free and the high fish predation (sub)populations (i.e. between the Pre-fish and High-fish subpopulation of Cousyn et al. (2001) and between Citadelpark and Blankaart population in De Meester (1996)) nor did we find differences between the high and reduced fish predation (sub)populations (i.e. contributions between High-fish and Reduced-fish subpopulation of Cousyn et al. (2001) and between Blankaart and Driehoeksvijver population of De Meester (1996)) (Fig. F8C,D).

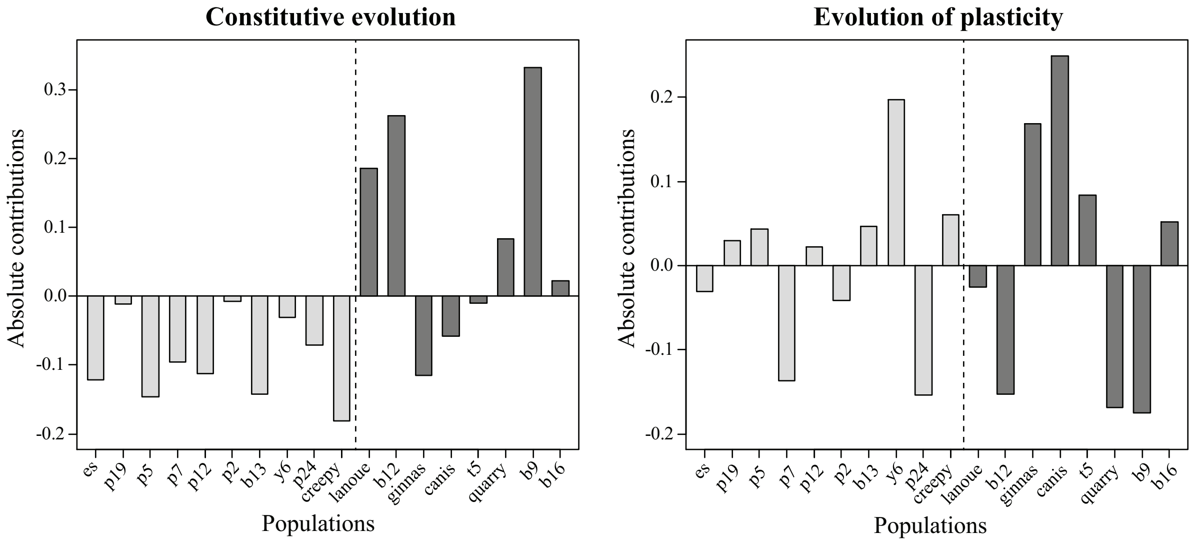


Figure F1: Absolute contributions of constitutive evolution (A) and evolution of plasticity (B) to trait shifts from an average predator-free *A. maculatum* population to each of the predator-present populations (light grey bars) and to trait shifts from an average predator-present *A. maculatum* population to each of the predator-free populations (dark grey bars).

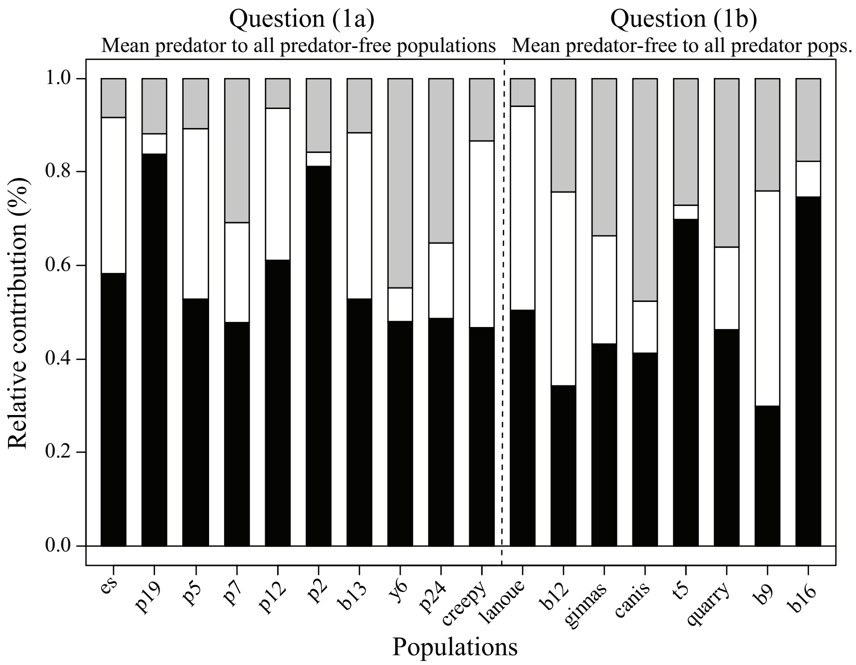


Figure F2: Relative contributions of plasticity (black), constitutive evolution (white) and evolution of plasticity (grey) to change in mean prey body size going from an average predator-present *A. maculatum* population to each of the predator-free populations (Question (1a)), and to change in mean prey body size going from an average predator-free *A. maculatum* population to each of the predator-present populations (Question (1b)).

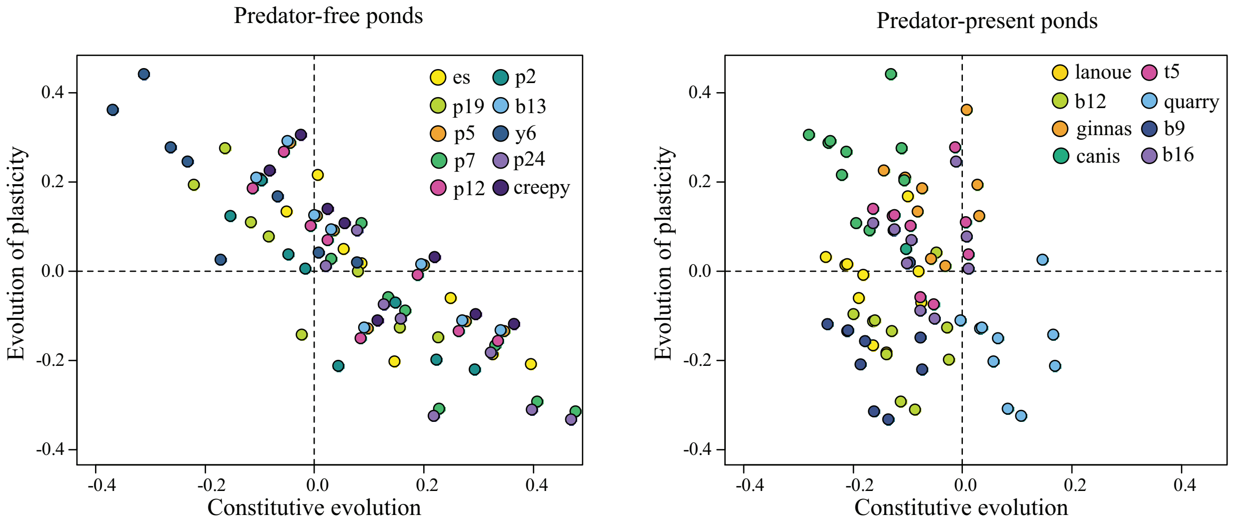


Figure F3: Absolute contributions of constitutive evolution and evolution of plasticity to trait shifts (A) from each of the predator-free *A. maculatum* populations to each of the predator-present populations, and (B) from each of the predator-present *A. maculatum* populations to each of the predator-free populations. Colors indicate calculated contributions using the same reference populations. For example, in (A) yellow dots reflect contributions of constitutive evolution and evolution of plasticity to shift in mean prey size using the *A. maculatum es* population as a reference to each of the 8 predator-present populations.

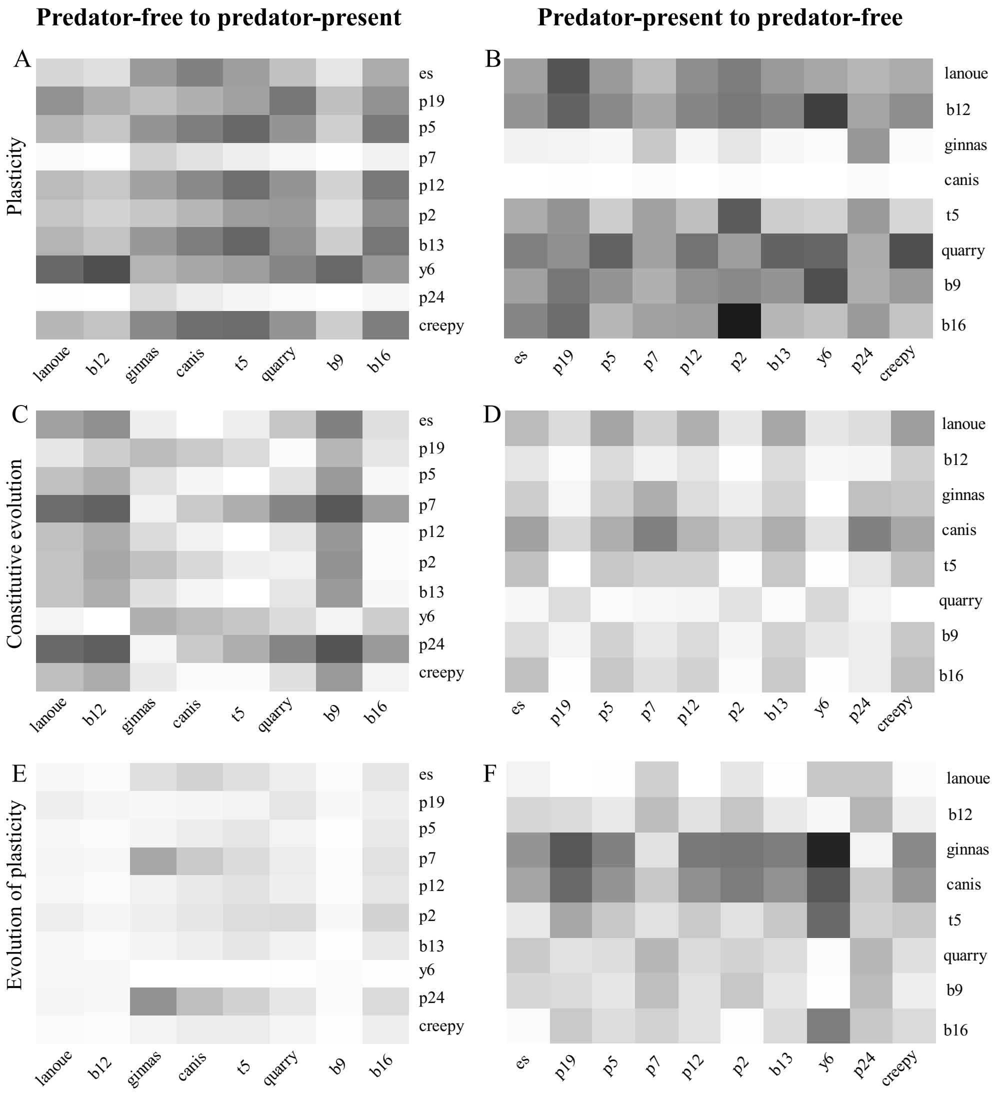


Figure F4: Heatmap representation of the relative contributions of plasticity (A, B), constitutive evolution (C, D) and evolution of plasticity (E, F) to change in prey body size going from each of the predator-free *A. maculatum* populations to each of the predator-present *A. maculatum* populations (Question (2a); A, C, E), and when going from each of the predator-present *A. maculatum* populations to each of the predator-free *A. maculatum* populations (Question (2b); B, D, F). Shades of grey are comparable across plots, and darker shades of grey indicate larger relative contributions, with black reflecting a contribution of 100%.

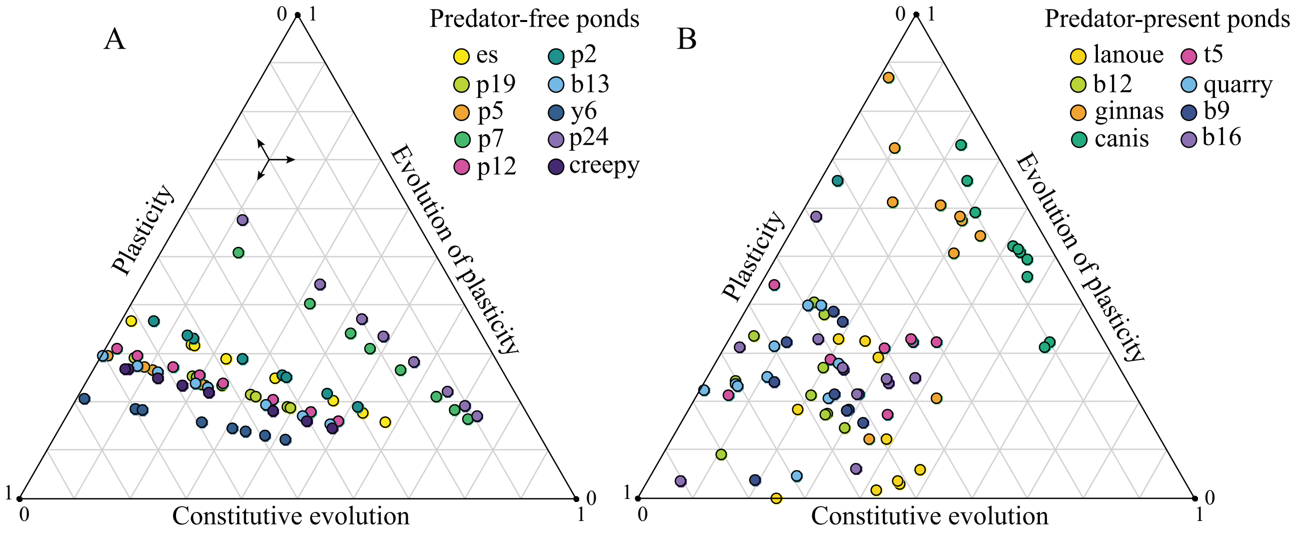


Figure F5: Alternative visualization of the results of Question (2a) and (2b) using triangle plots. Triangle plots show the relative contributions of plasticity, constitutive evolution and evolution of plasticity to change in prey body size going from each predator-free *A. maculatum* population to each of the predator-present populations (Question (2a); A), and when going from each of the predator-present *A. maculatum* populations to each of the predator-free populations (Question (2b); B). Each color represents the contributions of plasticity, constitutive evolution and evolution of plasticity obtained when using the same reference population. For example, in (A) yellow dots represent the contributions when going from the predator-free *A. maculatum es* population to each of the predator-present populations (for total overview see color code in figure legend). Arrows indicate how to read the triangle plot.

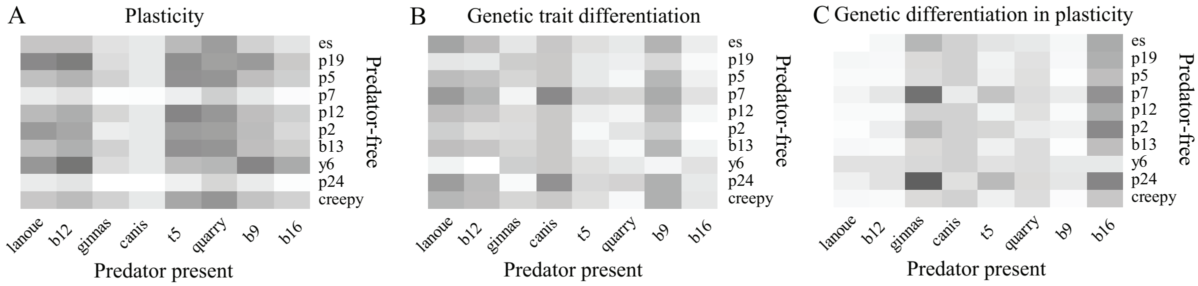


Figure F6: Heatmap representation of the relative contributions of (A) plasticity, (B) genetic trait differentiation and (C) genetic differentiation in plasticity to shifts in prey body mass between pairwise comparisons of *A. maculatum* populations originating from habitats where the predator *A. opacum* is either present or absent.

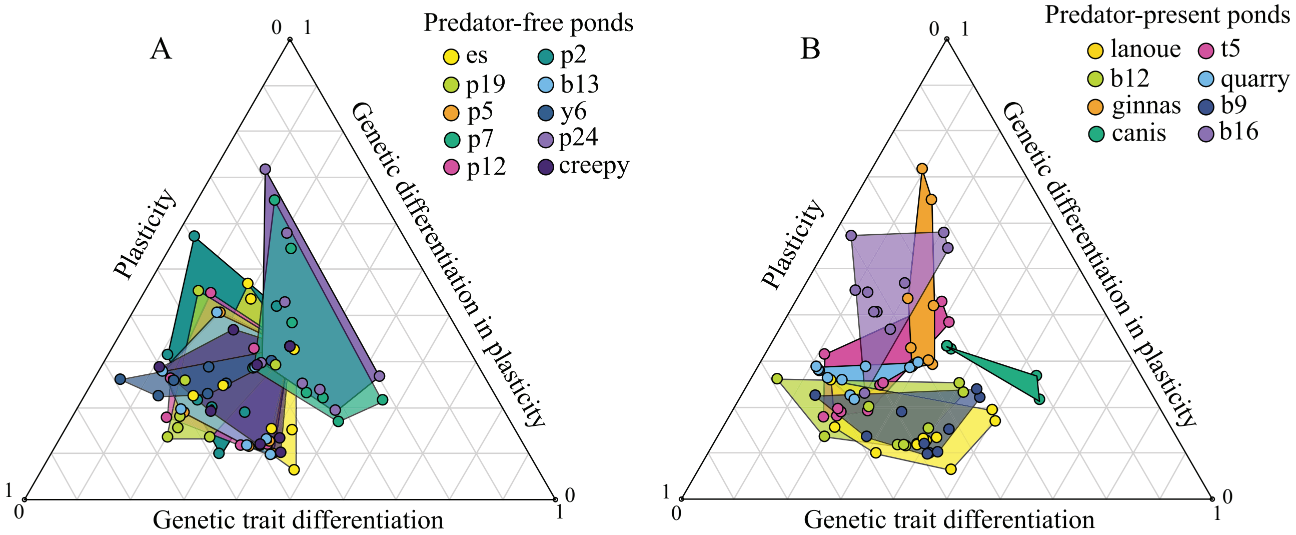


Figure F7: Alternative visualization of the relative contributions of plasticity, genetic trait differentiation and genetic differentiation in plasticity to shifts in prey body mass between pairwise comparisons of *A. maculatum* populations originating from habitats where the predator *A. opacum* is either present or absent using triangle plots. In (A) relative contributions are clustered by pairs that contain the same predator-free population. In (B) relative contributions are clustered by pairs that contain the same predator-present population. Colors refer to the respective population to which the clustering is done.

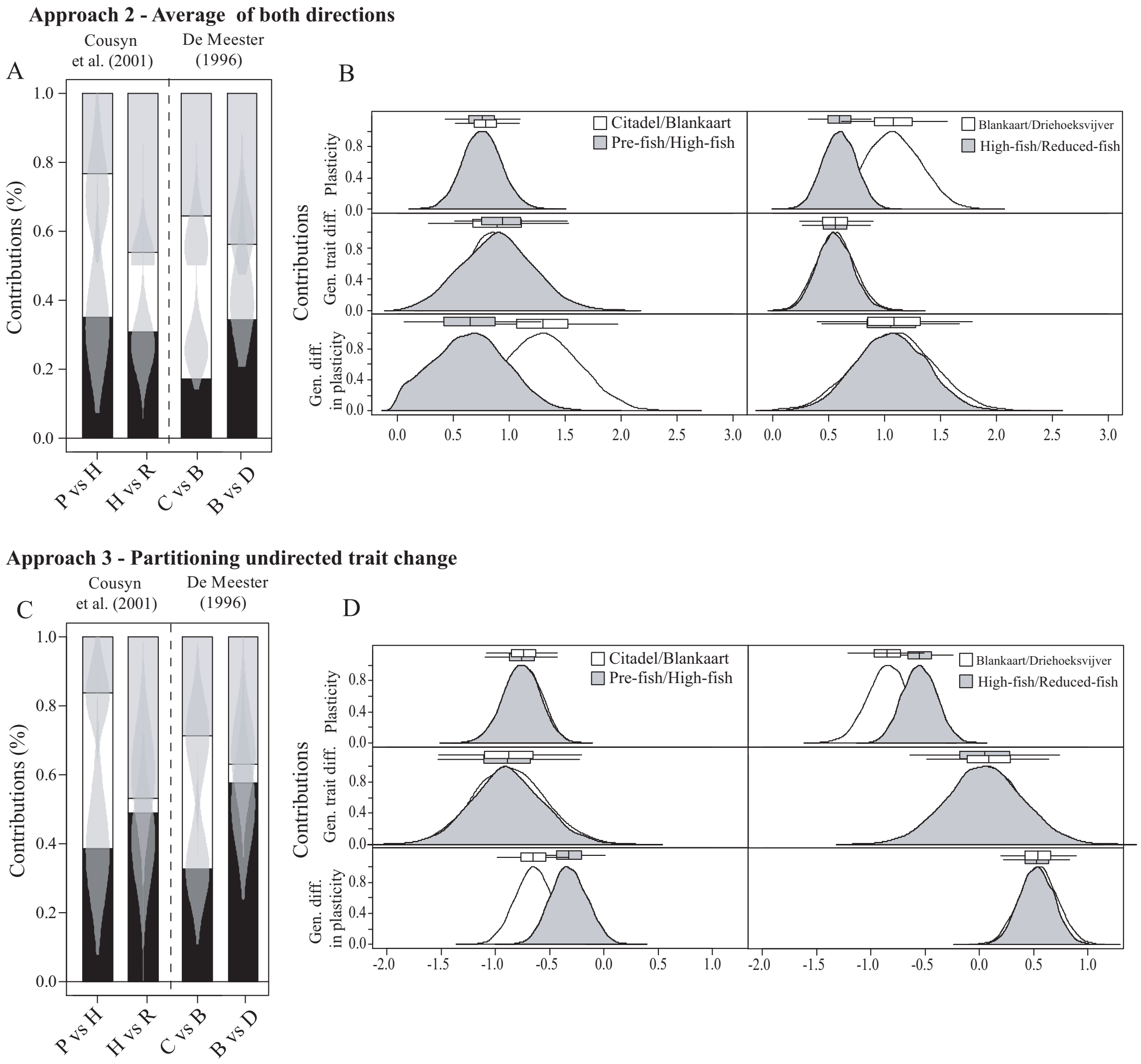


Figure F8: Relative and absolute contributions and density distributions of the obtained bootstrap values (shaded area) of plasticity (black), genetic trait differentiation (white) and genetic differentiation in plasticity (grey) to change in phototactic behaviour using (A-B) Approach 2 and (C-D) Approach 3 on the Cousyn et al. (2001) data between the Pre-fish and High-fish (P vs H) and between the High-fish and Reduced-fish (H vs R) *D. magna* subpopulations and on the De Meester (1996) data between the *D. magna* populations from Citadel and Blankaart (C vs B) and from Blankaart and Driehoeksvijver (B vs D). (B, D) display density distributions obtained from the bootstrapping for the absolute contributions of plasticity, genetic trait differentiation and genetic differentiation in plasticity between the Pre-fish and High-fish and between the High-fish and Reduced-fish subpopulation from the Cousyn et al. (2001) data, and between the Citadel and Blankaart and between the Blankaart and Driehoeksvijver from the De Meester (1996) data.
